# Deletion and overexpression of the scaffolding protein IQGAP1 promotes HCC

**DOI:** 10.1101/2020.05.29.124404

**Authors:** Evan R Delgado, Hanna L Erickson, Junyan Tao, Satdarshan P Monga, Andrew W Duncan, Sayeepriyadarshini Anakk

**Affiliations:** Department of Pathology, McGowan Institute for Regenerative Medicine, Pittsburgh Liver Research Center, University of Pittsburgh, Pittsburgh, PA; Department of Molecular and Integrative Physiology, Cancer Center at Illinois, University of Illinois at Urbana-Champaign, Urbana, IL 61801

## Abstract

IQ motif–containing GTPase-activating protein 1 (IQGAP1) is a ubiquitously expressed scaffolding protein that is overexpressed in a number of cancers, including liver cancer, and is associated with many pro-tumorigenic processes including cell proliferation, motility, and adhesion. IQGAP1 can integrate multiple signaling pathways and could be an effective anti-tumor target. Therefore, we examined the role for IQGAP1 in tumor initiation and promotion during liver carcinogenesis. Unexpectedly, we found that *Iqgap1*^*-/-*^ mice had a higher tumor burden than *Iqgap1*^*+/+*^ and *Iqgap1*^*+/-*^ mice following DEN-induced liver carcinogenesis. *Iqgap1*^*-/-*^ tumors as well as knocking down *IQGAP1* in hepatocellular carcinoma (HCC) cell lines resulted in increased MET activation and cellular proliferation. On the other hand, we uncovered IQGAP1 overexpression accelerates HCC development by YAP activation and subsequent NUAK2 expression. We demonstrate that increasing IQGAP1 expression *in vivo* does not alter β-catenin or MET activation. Taken together, we identify that both loss and gain of function of IQGAP1 promotes HCC development by two separate mechanisms in the liver. These results demonstrate that adequate amount of IQGAP1 is necessary to maintain a quiescent status of liver.

## Introduction

Liver cancer has a high mortality rate that is mainly attributed to lack of effective systemic therapies [1]. For hepatocellular carcinoma (HCC), the major form of primary liver cancer, a large portion of cases are diagnosed at advanced stages [2] and only two systemic therapies extend overall survival by a few months [1, 3]. Both modalities are multi-kinase inhibitors where Sorafenib targets the serine-threonine kinases Raf-1 and B-Raf, vascular endothelial growth factor receptors (VEGFR) 1-3, and platelet-derived growth factor receptor (PDGFR) [1] and Lenvatinib targets VEGFR 1-3, fibroblast growth factor receptors (FGFR) 1-4, PDGFRα, stem cell factor receptor (KIT), and rearranged during transfection (RET) [3]. Many other kinase inhibitors have failed to improve on this survival benefit in phase III trials [4-9]. It is clear that a better understanding of liver tumor biology is required to improve therapeutic strategies for HCC.

IQ motif–containing GTPase-activating protein 1 (IQGAP1) is a pleiotropic, multi-domain scaffolding protein that is overexpressed in many types of human cancer [10], including 60-80% of HCCs [11-14], and this overexpression is associated with worse clinical outcomes [13]. IQGAP1 interacts with pro-tumorigenic processes, including kinase signaling, cell proliferation, motility, and adhesion [10]. Furthermore, *in vivo* studies demonstrate that increased IQGAP1 expression can promote tumor growth, indicating that IQGAP1 could be an effective molecular target for HCC [15-17]. Yet, other studies revealed that deletion of IQGAP1 in cancer cells and/or stromal cells can also enhance tumorigenesis by modulating transforming growth factor (TGF) signaling [18, 19] and adherens junction stability [20].

To understand these contradictory studies, in this paper we carefully investigated the role for IQGAP1 in hepatic tumorigenesis by directly comparing IQGAP1-knockout (*Iqgap1*^*-/-*^) and IQGAP1-overexpression mouse models of HCC. Using the diethylnitrosamine (DEN) model of liver cancer, we show that deletion of both copies of *Iqgap1* increases the incidence and multiplicity of liver tumors. *Iqgap1*^*-/-*^ tumors exhibited elevated levels of the tyrosine kinase receptor MET, and this could directly contribute to increased proliferation. Further, we show that overexpression of IQGAP1 enhances rapid onset and progression of HCC. YAP1 activation driven by an increase in NUAK2 kinase expression contribute to this model. Thus, we show that both deletion and overexpression of IQGAP1 can promote the development of liver tumors in mice using two distinct molecular mechanisms. Our findings underscore that molecular expression of liver tumors should be considered when developing new therapies for HCC.

## Results

### Deletion of IQGAP1 promotes DEN-induced tumorigenesis

To study whether IQGAP1 is required for hepatic tumorigenesis, we used the DEN model of liver cancer, a gold standard for chemical carcinogenesis that effectively mimics the prolonged development of HCC in humans following a previously published protocol [21]. We treated male and female *Iqgap1*^*+/+*^, *Iqgap1*^*+/-*^, and *Iqgap1*^*-/-*^ mice with 5 mg/kg DEN via intraperitoneal injection at 12-15 days of age and assessed tumor burden 20- and 50-weeks post-treatment. Visible tumors were observed in 0/9 *Iqgap1*^*+/+*^, 1/3 *Iqgap1*^*+/-*^, and 0/3 *Iqgap1*^*-/-*^ female mice at the 50-week time point, which is consistent with previous reports that females are protected from developing HCC [22, 23].

Because female mice are protected from developing HCC, we performed all further analysis on male mice only. In the male mice, no macroscopic nodules were observed at 20 weeks (Supplemental Figure 1A), but microscopic lesions were observed in 3/9 *Iqgap1*^*+/+*^ (33.3%), 3/10 *Iqgap1*^*+/-*^ (30%), and 3/5 *Iqgap1*^*-/-*^ (60%) livers at that time point (Supplemental Figure 1B). Macroscopic nodules were observed at 50 weeks post DEN treatment in 9/13 *Iqgap1*^*+/+*^ (69.2%), 11/24 *Iqgap1*^*+/-*^ (45.8%), and 14/16 *Iqgap1*^*-/-*^ (87.5%) mice (Figure 1A-B). We found higher incidence (Figure 1B) and multiplicity (Figure 1C) of liver tumors in *Iqgap1*^*-/-*^ compared to *Iqgap1*^*+/-*^ mice, indicating that total loss of *Iqgap1* promotes tumorigenesis. However, *Iqgap1-*deletion did not affect maximum tumor size (Figure 1D). When we segregate a subset of mice with LW/BW >7% (Figure 1E), we found that *Iqgap1*^*-/-*^ livers were significantly larger than *Iqgap1*^*+/+*^ and *Iqgap1*^*+/-*^ livers. These data suggest that the tumor burden is higher in *Iqgap1*^*-/-*^ animals.

**Figure 1.**
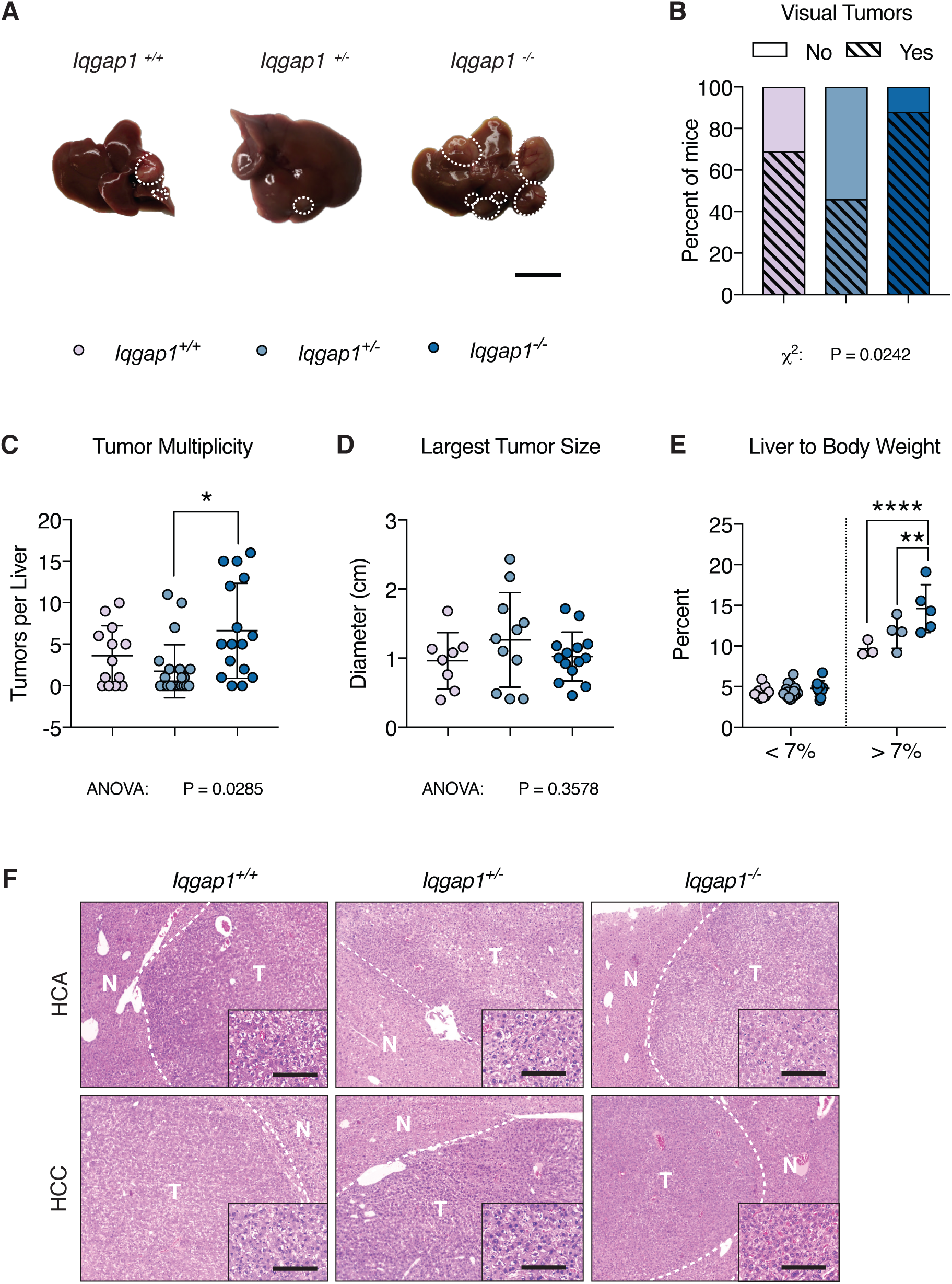
IQGAP1 deletion promotes liver tumor development. Male *Iqgap1*^*+/+*^ (n = 13), *Iqgap1*^*+/-*^ (n = 24), and *Iqgap1*^*-/-*^ (n = 16) mice were treated intraperitoneally with 5 mg/kg diethylnitrosamine (DEN) in sterile PBS at 12-15 days of age. At 1 year, tumor burden was assessed. (A) Representative photos of gross liver. Tumor nodules are indicated by a white dashed border. Scale bar is 1 cm. (B) Tumor incidence based on presence of visible liver nodules. (C) Tumor multiplicity was measured by counting the number of visible tumors per liver. (D) Tumor size was assessed by measuring the diameter of the largest visible tumor. (E) Liver weight normalized to body weight divided into low and high groups. (F) Representative H&E images of normal liver tissue, hepatocellular adenoma (HCA), and hepatocellular carcinoma (HCC) in tumor-bearing *Iqgap1*^*+/+*^, *Iqgap1*^*+/-*^, and *Iqgap1*^*-/-*^ mice (n = 9, 8, and 12 mice per group, respectively). Scale bar is 200 µm; 100 µm (inset). Values are displayed as mean ± SD. For tumor incidence, χ^2^ test was used to determine significance between all 3 groups. For tumor multiplicity and largest tumor size, one-way ANOVA with Bonferroni’s multiple comparison test was used to determine significance between groups. For liver weight and liver to body weight ratio, two-way ANOVA with Bonferroni’s multiple comparison test was used to determine significance between groups. Significance is indicated by * *P* < 0.05, ** *P* < 0.01, *** *P* < 0.001, **** *P* < 0.0001.

We previously showed that *Iqgap1*^*-/-*^ mice have lower gonadal adipose tissue weight [24]. Since body composition can both affect and be affected by tumor burden, we asked whether the increased tumor burden in *Iqgap1*^*-/-*^ animals corresponded to differences in body composition. Body weight was approximately 14% lower in *Iqgap1*^*-/-*^ animals (31.2 g) than both *Iqgap1*^*+/+*^ (36.4 g) and *Iqgap1*^*+/-*^ (37.0 g) mice at 50 weeks post-DEN treatment (Supplemental Figure 2A). The same trend was observed as early as 10 weeks of age, indicating that the effect of *Iqgap1-*deletion on body weight occurred prior to tumorigenesis (Supplemental Figure 2B). At this time point, no difference in white adipose tissue (WAT) or quadriceps muscle weight was observed (Supplemental Figure 2C-D). Levels of serum injury markers alanine aminotransferase (ALT), aspartate aminotransferase (AST), and total bilirubin were elevated but not different between groups (Supplemental Figure 2E-G). This suggests that the increased tumor burden in *Iqgap1*^*-/-*^ animals is likely not due to differences in body composition or increased liver injury.

**Figure 2.**
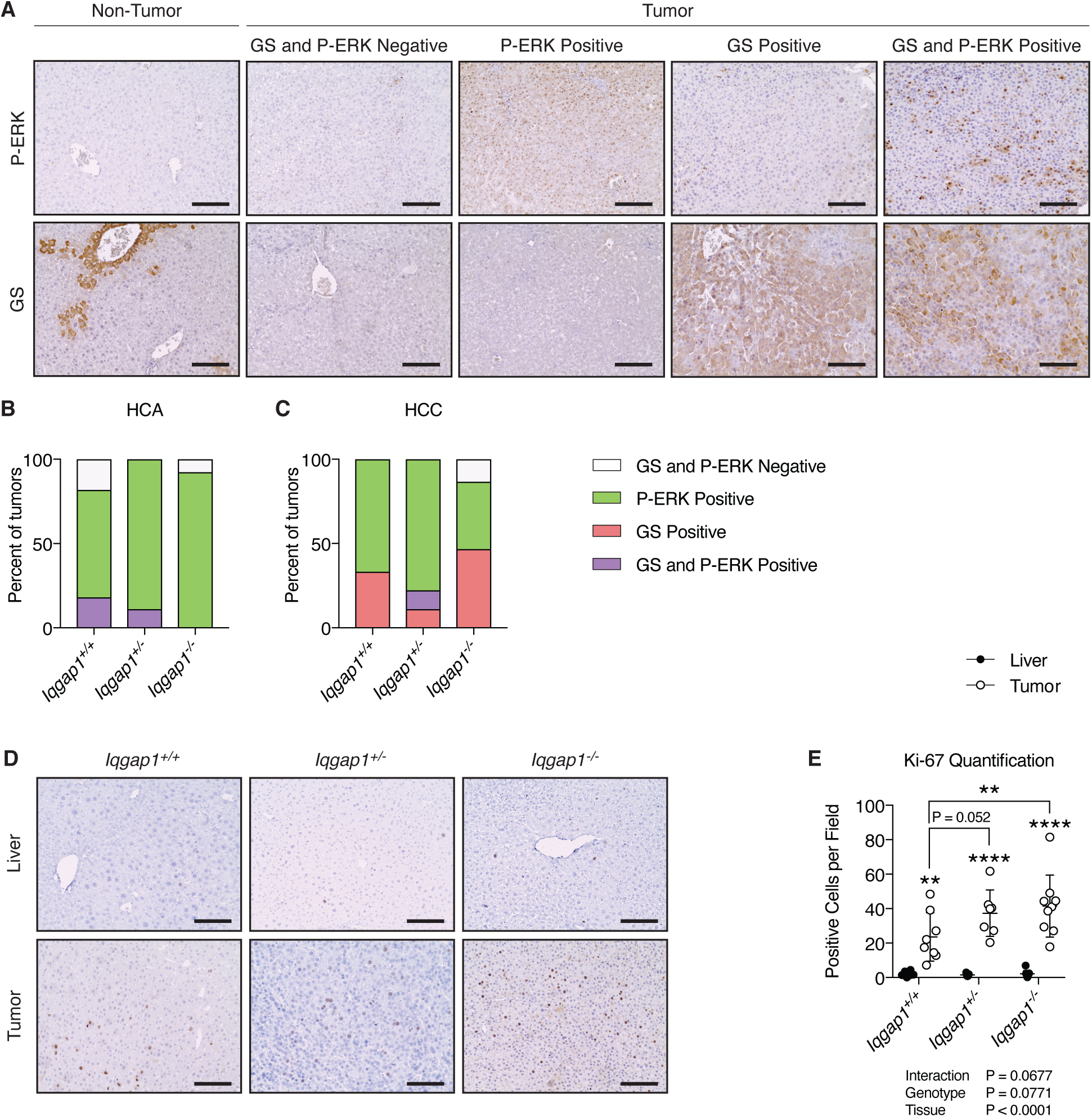
DEN-induced tumors are either P-ERK or GS positive. (A) Representative immunohistochemistry images of anti-P-ERK and anti-GS staining in normal liver tissue and tumor tissue classified as either GS and P-ERK negative, P-ERK positive, GS positive, or GS and P-ERK positive. Scale bar is 100 µm. (B) The percent of HCA tumors with each molecular classification (n = 12, 9, and 15 *Iqgap1*^*+/+*^, *Iqgap1*^*+/-*^, and *Iqgap1*^*-/-*^ tumors, respectively). (C) The percent of HCC tumors with each molecular classification (n = 11, 9, and 13 *Iqgap1*^*+/+*^, *Iqgap1*^*+/-*^, and *Iqgap1*^*-/-*^ tumors, respectively). (D) Representative images of anti-Ki-67 immunohistochemistry staining in liver and tumor tissue of *Iqgap1*^*+/+*^, *Iqgap1*^*+/-*^, and *Iqgap1*^*-/-*^ animals (n = 9, 8, and 12 mice per group, respectively). Scale bar is 100 µm. (E) Quantification of proliferating (Ki-67 positive) nuclei in liver and tumor tissue. Values are displayed as mean ± SD. Two-way paired ANOVA with Tukey’s multiple comparisons test was used to assess differences between groups. Significance is indicated with * *P* < 0.05, ** *P* < 0.01, *** *P* < 0.001, **** *P* < 0.0001.

We next confirmed that deletion of *Iqgap1* did not result in compensatory expression changes in homologs *Iqgap2* and *Iqgap3*. Of the three isoforms, IQGAP1 has the widest tissue distribution and is more frequently altered in cancer [25]. As expected, *Iqgap1* was induced in *Iqgap1*^*+/+*^ tumors relative to the surrounding healthy liver, and the *Iqgap1*^*+/-*^ mice exhibited an approximately 50% reduction in *Iqgap1* expression compared to controls (Supplemental Figure 3A). *Iqgap2* is more highly expressed in the control liver (*Iqgap1*^*+/+*^ liver Cq = 18-19) than *Iqgap1* (*Iqgap1*^*+/+*^ liver Cq = 23-24) but is decreased in tumor tissue (Supplemental Figure 3B). Whereas *Iqgap3* expression is low in the quiescent livers (*Iqgap1*^*+/+*^ liver Cq = 30-31), it is observed in proliferating cells [26, 27] and dramatically induced in tumor tissue (Supplemental Figure 3C). Notably, expression pattern of neither *Iqgap2* nor *Iqgap3* was altered by *Iqgap1-*deletion.

**Figure 3.**
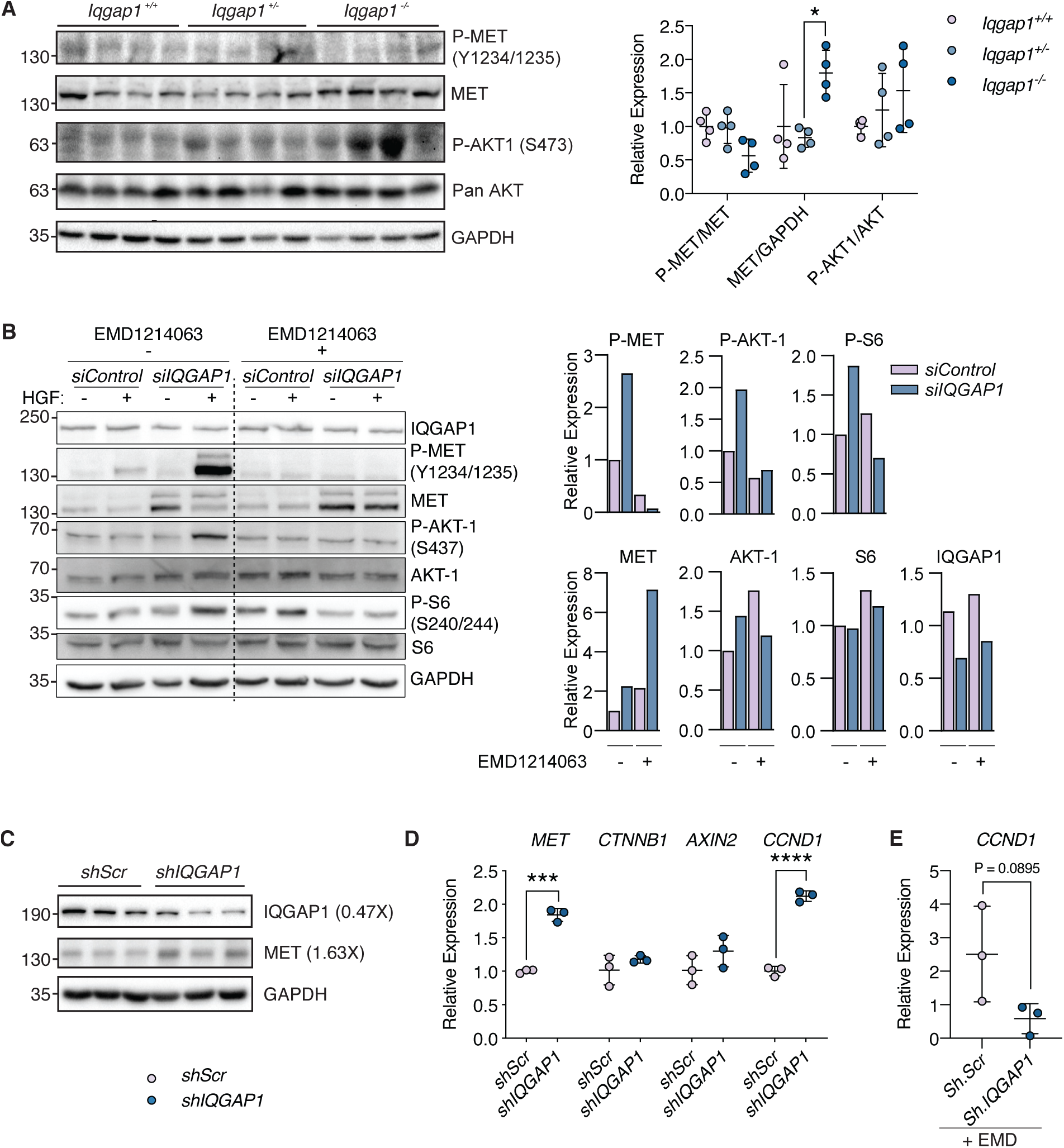
IQGAP1 loss enhances MET expression and activation. (A) Immunoblot of DEN-treated *Iqgap1*^*+/+*^, *Iqgap1*^*+/-*^, and *Iqgap1*^*-/-*^ tumor extracts. Each lane contains extracts from a single mouse. Quantification of immunoblots by densitometry provided at right. (B) Snu-449 HCC cells were transfected with either *Control* or *IQGAP1* siRNA for 48 hours. Cells were then serum starved overnight and treated with ±HGF (50 ng/µL) and ±EMD1214063 (10 nM) for 4 hours prior to harvest. Whole cell lysates were immunoblotted for phosphorylated and total forms of specific targets and normalized to GAPDH. Conditions are representative of 3 independent experiments pooled together. Quantification of immunoblots with respect to only HGF+ condition by densitometry shown at right. (C) Immunoblot of HepG2 cells infected with lentivirus expressing *shScrambled* (*shScr*) and *shIQGAP1*. (D-E) Gene expression of HepG2 cells infected with *shScr* and *shIQGAP1* (D) and treated with EMD1214063 (E) normalized to *Gapdh* expression. Values are displayed as mean ± SD. To compare 2 groups, Student’s t-test was performed. Two-way paired ANOVA with Tukey’s multiple comparisons test was used to assess differences between groups. Significance is indicated with * *P* < 0.05, ** *P* < 0.01, *** *P* < 0.001, **** *P* < 0.0001.

### Highly proliferative *Iqgap1*^*-/-*^ tumors are characterized by elevated GS or activated ERK

The incidence and multiplicity of tumors were higher in *Iqgap1*^*-/-*^ mice, so we asked whether the tumors differed in proliferation or pathology. The DEN model generates both hepatocellular adenoma (HCA), a benign tumor, and hepatocellular carcinoma (HCC), a malignant tumor [21]. These can be histologically differentiated as HCC has a thick trabecular structure, while HCA has cells arranged in thin trabeculae or in cords similar to normal hepatocytes with no vasculature. Since some of the livers had multiple tumors, we characterized each tumor separately. Both tumor types were observed in all three genotypes and some livers had both HCA and HCC tumors. (Figure 1F). HCA nodules were found in 11/23 *Iqgap1*^*+/+*^ (47.8%), 9/18 *Iqgap1*^*+/-*^ (50.0%), and 13/28 *Iqgap1*^*-/-*^ (46.4%) tumors while HCC nodules were found in 12/23 *Iqgap1*^*+/+*^ (52.1%), 9/18 *Iqgap1*^*+/-*^ (50%), and 15/28 *Iqgap1*^*-/-*^ (53.6%) tumors.

We next asked if there were any fundamental differences in the molecular characteristics of these tumors. Liver cancer can be divided into at least six molecular subtypes (G1-G6) depending on their gene expression pattern (Supplemental Table 1) [28]. We analyzed markers of proliferation (Supplemental Figure 3D-E), lipogenesis (Supplemental Figure 3F), angiogenesis (Supplemental Figure 3G), inflammation (Supplemental Figure 3H), gluconeogenesis (Supplemental Figure 3I), and β-catenin activity (Supplemental Figure 3J) that have been previously identified to correspond to these distinct molecular subtypes of human HCC [28, 29]. We found no difference in gene expression pattern between groups. The expression changes of 5/7 genes (*Rrm2, Tgfbr1, Fasn, Pepck, Angpt2*) aligned with G3 tumors, which are characterized by chromosomal instability, cell cycle activation, and *TP53* mutations [28].

DEN promotes hepatocellular carcinogenesis by inducing covalent DNA adducts in hepatocytes that can increase the mutation rate and thus the risk of uncontrolled growth. There are 4 codons known to be frequently altered in DEN-induced tumors (BRAF K584, BRAF V637, EGFR F254, and HRAS Q61) [30, 31], and we assessed whether the prevalence of these DNA mutations varied among the groups as a marker for overall mutation rate. Of these, only BRAFV637 showed considerable percent mutation (Supplemental figure 4A-D). However, there was no difference in the amount mutated between groups. Furthermore, we assessed the initial proliferative response to DEN treatment at 24 hours after injection since an impaired response to injury can drive tumor growth. We found no difference in proliferation rate between groups (Supplemental figure 4E-F). Together, these data show that *Iqgap1* deletion does not affect mutation rate or the initial proliferative response to DEN.

**Figure 4.**
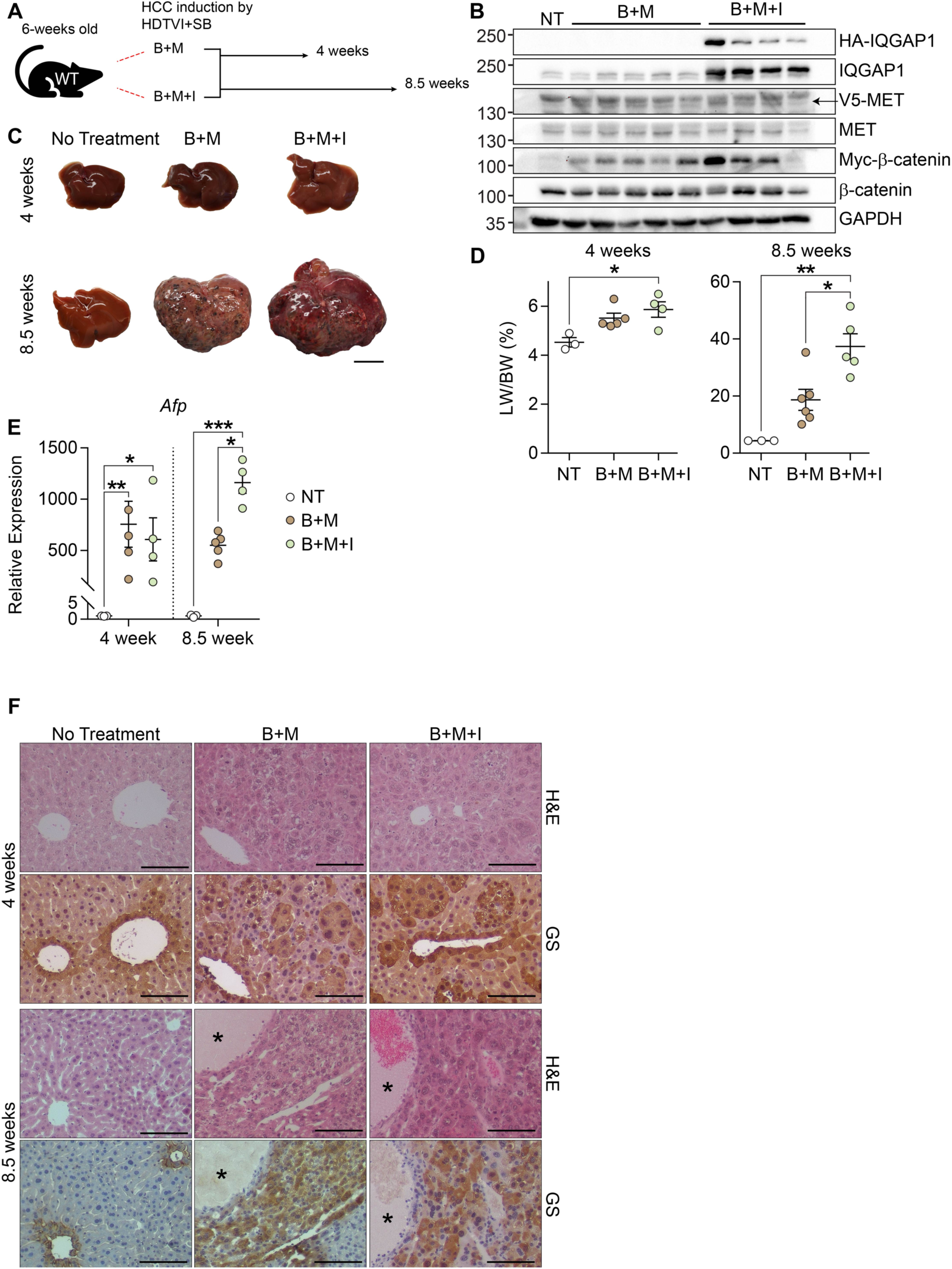
IQGAP1 overexpression increases HCC development in B+M transposon system. (A) Experimental design for Sleeping Beauty Transposase groups injected with either B+M or B+M+I and harvested after 4 or 8.5 weeks. (B) Whole liver protein lysates isolated after 4 weeks and analyzed by western blot to detect epitope tagged (HA, V5 or Myc) and total IQGAP1, MET, and β-catenin; normalized to GAPDH. (C) Representative livers from non-treated, B+M, and B+M+I groups. Macroscopic disease is visible as black or white lesions. (D) Liver weight to body weight (LW/BW) ratio. (E) *Afp* expression corrected for *Gapdh*. (F) Serial liver sections from 4-week and 8.5-week mice stained with H&E and GS (brown) to identify β-catenin driven HCCs. Necrotic regions are marked by asterisks. Graphs show mean ± SEM and dots represent individual mice. Two-way paired ANOVA with Tukey’s multiple comparisons test was used to determine significance between groups. Significance is indicated with * *P* < 0.05, ** *P* < 0.01, *** *P* <0.001. Gross morphology scale bar (C) is 2.5 cm and histology scale (F) bars are 100 µm.

DEN-induced tumors are frequently driven by mutations in *Hras* and, to a lesser extent, mutations in *Ctnnb1* [32, 33]. We examined whether deletion of IQGAP1 altered the activation of these pathways by performing immunohistochemistry for phosphorylated ERK (P-ERK) and glutamine synthetase (GS). We divided tumors into four categories: GS and P-ERK negative, P-ERK positive, GS positive, and both P-ERK and GS positive (Figure 2A). IQGAP1 deletion did not affect the zonal expression pattern of GS in liver tissue. *Hras* and *Ctnnb1* mutations are typically mutually exclusive [34], which is consistent with our finding that a low number of tumors were both P-ERK and GS positive. HCAs were primarily P-ERK positive, with 9/11 *Iqgap1*^*+/+*^ (81.2%), 9/9 *Iqgap1*^*+/-*^ (100.0%), and 12/13 *Iqgap1*^*-/-*^ (92.3%) tumors shows positive staining (Figure 2B). On the other hand, a substantial proportion of HCCs had increased GS-positive staining (4/12 *Iqgap1*^*+/+*^ (33.3%), 2/9 *Iqgap1*^*+/-*^ (22.2%), and 7/15 *Iqgap1*^*-/-*^ (46.7%)) (Figure 2C).

Then we investigated if the increased tumor burden is associated with increased proliferation. Ki-67, a marker of non-quiescent cells, staining revealed low levels of positive cells in healthy liver tissue from tumor-bearing mice, regardless of genotype (Figure 2D-E). However, there was a stark increase in Ki-67 positive cells in tumors from *Iqgap1*^*-/-*^ mice compared to *Iqgap1*^*+/+*^ mice (Figure 2D-E). We checked markers of epithelial-mesenchymal transition in these tumors and found no significant difference in MMP2, E-cadherin, N-cadherin, or CDC42 protein expression between groups (Supplemental Figure 5). This indicates that *Iqgap1*-deletion does not alter the molecular characteristics of the tumors, but it does increase their proliferation.

**Figure 5.**
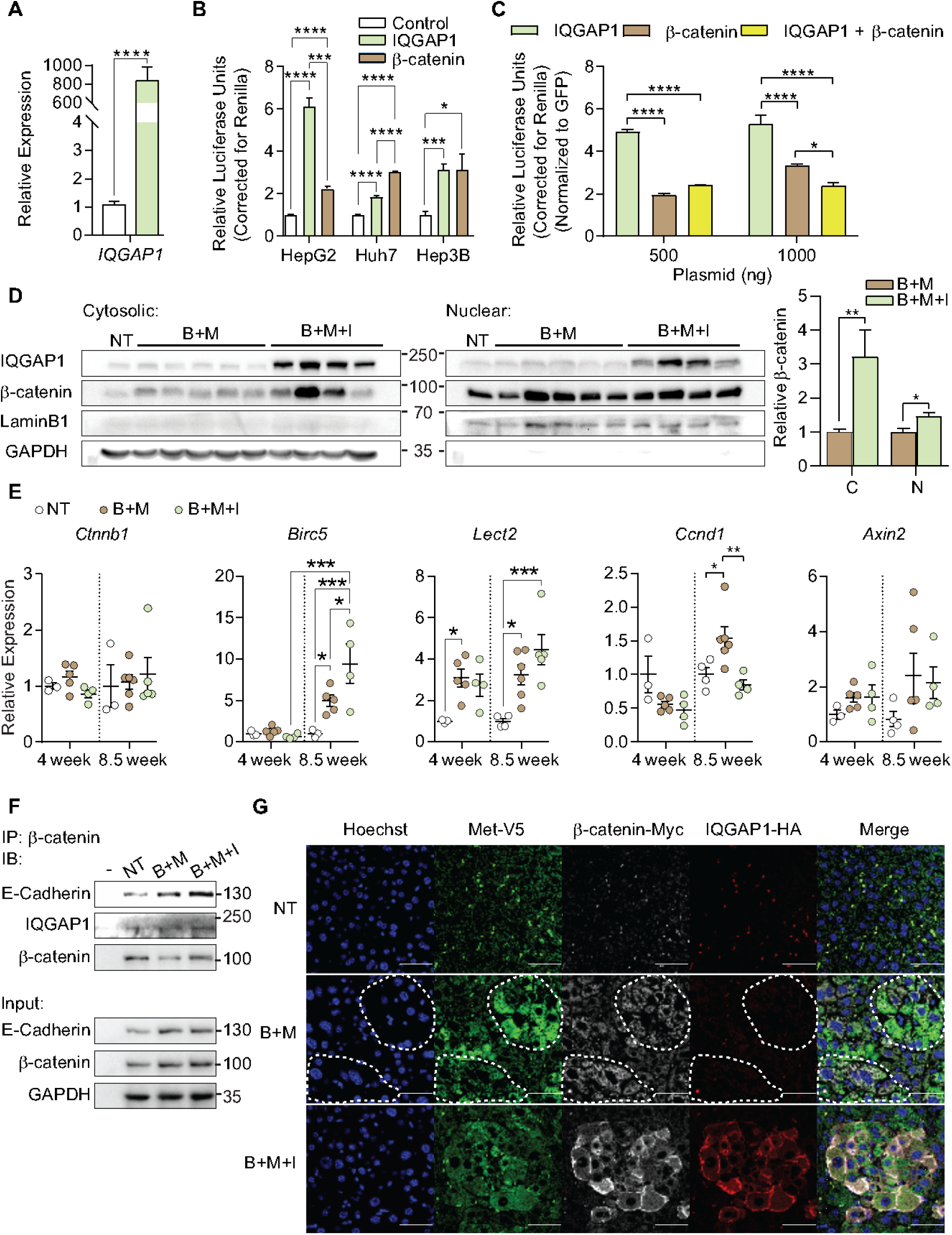
IQGAP1 overexpression does not induce Wnt/β-catenin signaling *in vivo*. (A) HepG2 cells were transfected with GFP-control or IQGAP1 constructs for 72 hours. Expression for *IQGAP1* was normalized to *B2M*. (B) HepG2, Huh7 and Hep3B cell lines were transfected with control, IQGAP1, or β-catenin expression constructs for 72 hours. Wnt/β-catenin activity was measured by the TOPFlash luciferase reporter assay and corrected for Renilla luciferase. (C) HepG2 cells were transfected with GFP, IQGAP1, or β-catenin expression constructs for 72 hours. Additionally, cells were also transfected with IQGAP1 and β-catenin constructs together. Wnt/β-catenin activity was measured and analyzed as previously described. (D) Cytosolic and nuclear protein from whole livers (4 week NT, B+M and B+M+I) were analyzed for IQGAP1 and β-catenin. Cytosolic protein was normalized to GAPDH and nuclear protein normalized to LaminB1. GAPDH and LaminB1 show purity of cytosolic or nuclear fractions, respectively. (E) Expression of *Ctnnb1* and Wnt/β-catenin target genes *Birc5, Lect2, Ccnd1* and *Axin2* in whole livers from 4- and 8.5-week samples, normalized to *Gapdh*. (F) Whole liver lysates from NT n=3, B+M n=5, or B+M+I n=4 were pooled and immunoprecipitated (IP) for β-catenin then immunoblotted (IB) for E-Cadherin, IQGAP1 or β-catenin. Sample inputs were probed for E-Cadherin, β-catenin or GAPDH to demonstrate equal amounts of protein from each group were used for IPs. (G) Liver sections from mice under the HDTVI model for 4 weeks stained for V5 (green), HA (red) or myc (white) to identify nodules by immunofluorescence expressing MET, IQGAP1 and β-catenin, respectively. Tumors in B+M condition are demarcated by white dashed line. Scale bar is 50 µm. Graphs show mean ± SEM and dots represent individual mice. For (E), two-way paired ANOVA with Tukey’s multiple comparisons test and all others compared via Student’s t-Test to determine significance. Significance is indicated with * *P* < 0.05, ** *P* < 0.01, *** *P* < 0.001, **** *P* < 0.0001.

### Loss of IQGAP1 results in dysregulated MET expression and activation

To identify the mechanism underlying tumorigenesis in *Iqgap1*^*-/-*^ DEN-mediated HCC model, we first examined the Wnt pathway. At the RNA level, *Ctnnb1* and its canonical target genes *Birc5*, and to a lesser extent *Axin2* and *Lect2* were induced in tumor tissue, regardless of genotype (Supplemental Figure 6A-D). Further, knock down of *IQGAP1* in HepG2 cells, Huh7, and Snu-449 cells revealed no change in β-catenin activity, as determined by TOPFlash luciferase reporter assay (Supplemental figure 6E-F). Also, DEN carcinogenesis resulted in GS positive nodules across all cohorts, suggesting that *Iqgap1* loss does not exacerbate Wnt signaling.

**Figure 6.**
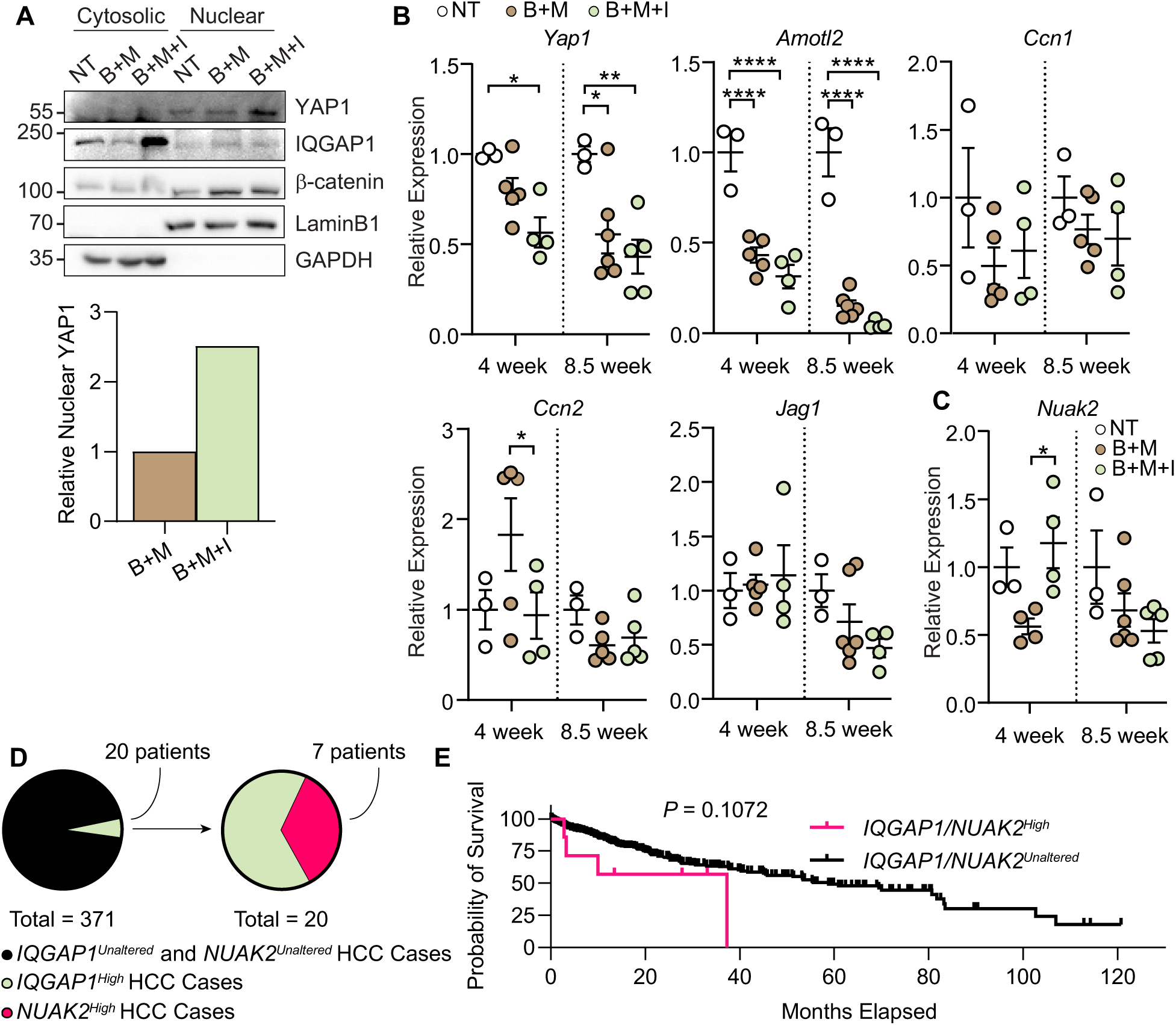
IQGAP1 overexpression drives HCC pathogenesis by inducing Hippo/YAP signaling *in vivo*. (A) Cytosolic and nuclear protein from whole livers (4 week pooled NT n=3, B+M n=5 and B+M+I n=4) were analyzed for YAP1, IQGAP1 and β-catenin. Cytosolic protein was normalized to GAPDH and nuclear protein normalized to LaminB1. GAPDH and LaminB1 show purity of cytosolic or nuclear fractions, respectively. (B) Expression of *Yap1* and Hippo/YAP target genes *Amotl2, Ccn1, Ccn2*, and *Jag1* in whole livers from 4- and 8.5-week samples, normalized to *Gapdh*. (C) Expression of *Nuak2* in whole livers from 4 and 8.5-week NT, B+M and B+M+I samples, normalized to *Gapdh*. (D) Pie charts demonstrating the distribution of HCC cases with *IQGAP1*^*High*^ expression from the TCGA cohort. Distribution further breaks down the number of cases with *IQGAP1*^*High*^ that also contain *NUAK2*^*High*^ expression. (E) Overall survival for subset of patients with *IQGAP1*^*High*^/*NUAK2*^*High*^ expression compared to *IQGAP1*^*Unaltered*^/*NUAK2*^*Unaltered*^ patients. Graphs show mean ± SEM and dots represent individual mice. For (B), two-way paired ANOVA with Tukey’s multiple comparisons test and for (C) Log-rank (Mantel-Cox) test was used to determine significance for survival curve and Student’s t-Test was used everywhere else. Significance is indicated with * *P* < 0.05, ** *P* < 0.01, **** *P* < 0.0001.

Since DEN-induced *Iqgap1*^*-/-*^ tumors display either activated P-ERK or GS expression, we investigated tyrosine kinase receptor MET, which can activate both Ras and β-catenin signaling pathways [35-38]. We measured MET expression and activation, marked by the phosphorylation of tyrosine residues 1234/1235 of MET (Y1234/1235) [38]. Although Y1234/1235 phospho-MET (P-MET) levels remained unchanged between the groups, total MET levels were elevated in *Iqgap1*^-/-^ tissues after DEN (Figure 3A). Phosphorylated AKT-1 (P-AKT-1) S473, a target downstream of MET signaling, also trended higher in *Iqgap1*^*-/-*^ tissues, which is consistent with a pattern of MET induction (Figure 3A). Next, we examined if *IQGAP1* knock down could directly regulate MET signaling *in vitro* using HepG2 and Snu449 cells. Knock down of IQGAP1 in cell lines was sufficient to increase MET expression (Figure 3B). Further, upon HGF stimulation IQGAP1 knock down cells showed increased phosphorylation of MET, AKT-1 and S6, suggesting that loss of *IQGAP1* renders cells highly sensitive to MET pathway activation (Figure 3B). In addition, treatment of cells with the MET small molecule inhibitor EMD1214063 blocks MET, AKT-1 and S6 phosphorylation after *IQGAP1* loss and HGF stimulation (Figure 3B). MET protein and RNA expression were also increased in HepG2 cells treated with shRNA for IQGAP1 (Figure 3C-D), but expression of β-catenin or its target *Axin1* expression were not affected (Figure 3D). To determine whether the increased MET expression correlated with increased proliferation, we also checked the expression of *CCND1*, which was 2- fold higher with *IQGAP1* knock down (Figure 3D). We can also block elevated *CCND1* following *IQGAP1* loss using EMD1214063 (Figure 3E). Together, the data indicate that loss of IQGAP1 results in increased MET signaling and proliferation, which could facilitate HCC oncogenesis.

### IQGAP1 overexpression exacerbates HCC carcinogenesis in the β-catenin/MET model

Since IQGAP1 expression is enhanced in the majority of HCC cases [11, 12], we asked whether overexpression of IQGAP1 was sufficient to promote tumor growth. To do this, we turned to a well-established HCC model that uses hydrodynamic tail vein injection (HDTVI) with the Sleeping Beauty (SB) transposase (referred hereafter as the “transposon system”). Briefly, the transposon system can induce HCC in wild-type (WT) adult mice by the forced expression of human activated (S45Y) β-catenin + MET (B+M) [39]. The S45Y mutation in β-catenin results in reduced degradation, enhanced nuclear translocation and activation of canonical Wnt signaling; the MET construct increases MET expression and downstream signaling events [39]. Simultaneous expression of B+M induces microscopic lesions visible by 2 weeks and macroscopic HCC within 6-9 weeks, which are 69% genetically similar to human HCC [39].

Using the transposon system, we overexpressed epitope tagged B+M with or without simultaneous expression of epitope tagged human IQGAP1 (B+M+I) in WT mice and harvested livers after 4 or 8.5 weeks (Figure 4A). As early as 4 weeks, tagged β-catenin and MET were expressed in liver lysates in B+M and B+M+I groups, compared to non-treated (NT) controls, and tagged IQGAP1 was expressed only in the B+M+I group (Figure 4B). We also checked expression of *Iqgap* family members and found that *Iqgap2* was unchanged while *Iqgap3* was elevated in both B+M and B+M+I livers compared to NT controls (Supplemental Figure 7A-B). Upregulation of *Iqgap3* in tumor tissues was also seen in DEN-induced tumors (Supplemental Figure 3C), again suggesting that *Iqgap3* induction marks cell proliferation.

**Figure 7.**
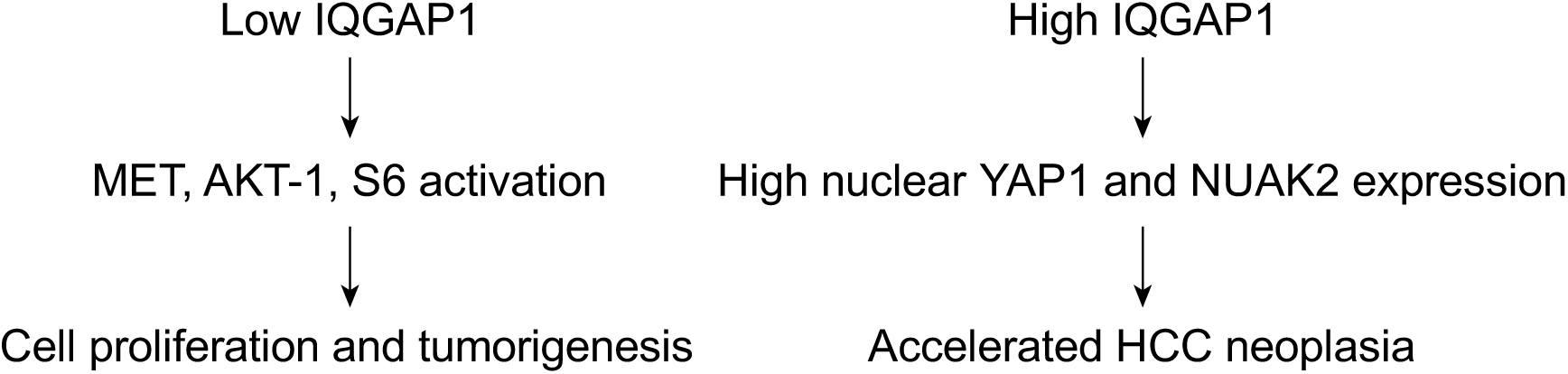
Summary of findings. Using two models of liver tumors, diethylnitrosamine, which produces both β-catenin and MAPK positive tumors, and a sleeping beauty hydrodynamic tail vein injection model overexpressing β-catenin and MET, we show that both overexpression and deletion of IQGAP1 promotes cell proliferation and increased tumor burden via enhanced Hippo signaling. IQGAP1 deletion results in increased MET activation and functions via AKT-1 and S6 signaling resulting in enhanced cell proliferation and tumor growth. IQGAP1 overexpression results in increased *NUAK2* expression permitting increased YAP1 nuclear translocation which promotes accelerated HCC neoplasia.

Macroscopic disease was not evident after 4 weeks, but at 8.5 weeks, there were visible macroscopic tumor nodules in both the B+M and B+M+I groups (Figure 4C). The LW/BW ratio (Figure 4D) was equivalent after 4 weeks but was 2-fold higher in the B+M+I group at 8.5 weeks compared to controls. Alpha-fetoprotein, a marker of highly aggressive HCC [40-42], was highest in the B+M+I group at 8.5 weeks (Figure 4E). Microscopic tumor nodules were present at each of the time points (Figure 4F). Tumors in B+M and B+M+I groups showed typical HCC morphology, visible by H&E staining, which contrasts with the mix of tumor types seen in the DEN model (Figure 1F). As expected for tumors induced by activated β-catenin, HCC nodules were enriched for GS expression in both groups and at all time points. In addition, we measured the expression of genes that differentiate between G1-G6 molecular subtypes, as described in Supplemental Table 1, and found changes in 4/7 genes (*Glul, Tgfbr1, Fasn*, and *Crp*) in B+M and B+M+I tissues are representative of G5/G6 tumors (Supplemental Figure 7C-I). Although *Crp* expression was lower in B+M+I tissues compared to B+M, we largely did not find differences in expression patterns between B+M and B+M+I samples. HCC tumors with activating *CTNNB1* mutations typically fall within the G5/G6 molecular subtypes, which are characterized by low cell proliferation, chromosomal stability, and a lack of inflammatory infiltrates [28, 29]. Notably, HCC nodules at 8.5 weeks were advanced and highly necrotic (Figure 4F). Thus, IQGAP1 overexpression in the B+M tumor model accelerates HCC tumor expansion.

### Overexpression of IQGAP1 does not accelerate HCC formation via enhanced MET or Wnt/β-catenin signaling

We sought to determine the mechanism by which IQGAP1 overexpression accelerates HCC formation in the transposon model. Deletion of IQGAP1 is associated with increased MET expression and activation (Figure 3), so we asked if IQGAP1 overexpression *in vivo* also impacts MET. MET is known to activate pro-tumorigenic pathways including MEK/ERK, PI3K/AKT/mTOR, and others, resulting in increased protein phosphorylation [38]. Therefore, we assessed P-MET Y1234/1235 in B+M and B+M+I livers (Supplemental Figure 8). Total MET expression was equivalent between groups, and phosphorylated MET at Y1234/1235 was 2-fold higher in B+M+I (Supplemental Figure 8). However, downstream targets of MET activation (AKT-1, mTOR, and STAT3) were unchanged (Supplemental Figure 8), suggesting that the MET pathway is not the driving mechanism of HCC in this model.

Multiple studies have shown that IQGAP1 enhances Wnt/β-catenin signaling *in vitro* [14, 43]. To verify this, we overexpressed IQGAP1 in cell culture (Figure 5A) and found 2 to 6-fold increase in Wnt/β-catenin activity upon IQGAP1 overexpression as measured by the TOPFlash reporter assay (Figure 5B). This induction was comparable to reporter activation achieved by an activated form of β-catenin alone. Next, we tested whether co-expression of IQGAP1 and β-catenin could increase reporter activity in an additive or synergistic manner. However, simultaneous expression of IQGAP1 and β-catenin did not change to β-catenin activity when compared to β-catenin alone (Figure 5C). Collectively, the data demonstrate that IQGAP1 overexpression can increase β-catenin activity *in vitro*, but this mechanism is more complex since overexpressing IQGAP1 and β-catenin together does not further enhance Wnt/β-catenin signaling.

IQGAP1 is reported to assist translocation of β-catenin from the cytoplasm to the nucleus *in vitro*, resulting in enhanced Wnt/β-catenin signaling [14]. Therefore, to directly test if IQGAP1 enhances β-catenin transcriptional activity *in vivo*, we investigated changes in β-catenin subcellular localization after IQGAP1 overexpression by isolating cytoplasmic and nuclear fractions from whole liver samples of B+M or B+M+I mice. As expected, IQGAP1 was elevated in the cytosol of B+M+I livers, compared to B+M (Figure 5D). This correlated with a striking 3-fold increase in cytosolic β-catenin (Figure 5D). In the nuclear fraction, a high level of IQGAP1 was found in B+M+I livers, demonstrating IQGAP1 translocation into the nucleus. Similarly, β-catenin levels showed modest but significant increase in the nucleus, consistent with previous findings (Figure 5D).

Since we and others have found increased IQGAP1 expression enhances β-catenin’s transcriptional activity *in vitro* (Figures 5B-C) [14, 43], we investigated if IQGAP1-mediated tumor promotion increases Wnt/β-catenin signaling *in vivo* by examining the expression of several canonical Wnt/β-catenin target genes. First, we checked β-catenin mRNA expression and, similar to the protein expression reported in Figure 4B, found no difference between B+M and B+M+I groups (Figure 5E). Next, after 4 weeks there was no significant difference in expression of canonical β-catenin targets *Birc5, Lect2, Ccdn1* or *Axin2* between B+M and B+M+I groups (Figure 5E). However, after 8.5 weeks, expression of target genes was highly variable. For example, *Birc5* was 2-fold higher in B+M+I livers compared to B+M livers; *Lect2* and *Glul* were significantly induced in both B+M+I and B+M livers; *Ccdn1* expression was induced in the B+M condition but not in B+M+I livers; and *Axin2* was unchanged. Thus, since *Birc5* expression is not exclusively controlled by Wnt/β-catenin signaling, and mRNA expression of *Ctnnb1, Lect2, Ccdn1, Axin2* and *Glul* (Supplemental Figure 6I) are similar between B+M and B+M+I groups, we concluded that IQGAP1 overexpression has a limited impact on Wnt/β-catenin signaling *in vivo*.

IQGAP1 is known to regulate the physical interaction between β-catenin and E-Cadherin at the cell membrane *in vitro* [44, 45]. We, therefore, investigated if IQGAP1 overexpression enhances β-catenin-E-Cadherin complexes *in vivo*. Using whole liver lysates from B+M or B+M+I mice, we co-immunoprecipitated β-catenin and probed for E-Cadherin and found IQGAP1 overexpression enhances β-catenin-E-Cadherin interactions (Figure 5F). We also stained B+M and B+M+I tissues for tagged β-catenin, MET and IQGAP1 and found that β-catenin and IQGAP1 co-localize at the cell membrane in B+M+I tissues (Figure 5G). These data consistently demonstrate that even though the B+M+I model shows increased tumor burden compared to the B+M model, enhanced Wnt/β-catenin signaling via IQGAP1 overexpression *in vivo* is not causing this result.

### IQGAP1 overexpression drives HCC via Hippo/YAP signaling

Our data indicate IQGAP1 does not enhance Wnt/β-catenin or MET signaling in the B+M model. Since we found elevated mRNA expression of *Birc5* and *Glul* (Figure 5E), which are also known targets of Hippo pathway [46, 47], we investigated its role in promoting tumorigenesis in the B+M+I model. Activation of Hippo kinases results in cytoplasmic retention of Yes-associated protein1 (YAP1) and thus inhibiting cellular proliferation. IQGAP1 has been shown to regulate YAP1 levels and activity [48, 49]. Therefore, we examined the cytoplasmic to nuclear translocation of YAP1 when IQGAP1 is overexpressed in the B+M model. We found that nuclear YAP1 (2.5-fold) was increased in the B+M+I samples, compared to B+M (Figure 6A). Next, we examined the mRNA expression of *Yap1* and its target genes (*Amotl2, Ccn1, Ccn2* and *Jag1*) and found that their expression largely remained unchanged between B+M and B+M+I groups except for *Ccn2* (Figure 6B). This data is consistent with recent findings that reveal *Amotl2, Ccn1, Ccn2* and *Jag1* have minimal functional role in HCC pathogenesis *in vivo* [50-55] even though they are important in tracking hepatoblastoma progression [56-59].

More recently, YAP1-driven HCC tumorigenesis was shown to be mediated by NUAK2 kinase, which was also shown to positively regulate YAP1 activity in a feed forward manner [60, 61]. Interestingly, we found *Nuak2* expression is maintained when IQGAP1 is overexpressed in the B+M model at the 4-week time-point compared to B+M alone where *Nuak2* is derepressed (Figure 6C). However, this regulation is not observed at the 8.5-week time-point (Figure 6C), which suggests that IQGAP1-YAP1 regulation may be lost in late stage of tumorigenesis. Consistently, elevated YAP1 activity is known as an early oncogenic event in HCC [62] and data from Figure 6A and C suggest that IQGAP1 may stabilize YAP1 activity by promoting NUAK2 expression at early stages of HCC oncogenesis. To further understand the clinical implications of the IQGAP1-NUAK2 axis in HCCs, using The Cancer Genome Atlas (TCGA), we identified that 35% of patients with *IQGAP1*^*High*^ mRNA expression also had *NUAK2*^*High*^ expression (Figure 6D, n=371). Patients with *IQGAP1*^*High*^/*NUAK2*^*High*^ expression exhibited a poor overall survival trend (Figure 6E) and their transcriptomic signature revealed activation of multiple pro-growth and pro-survival signaling pathways compared to other HCC patients (Supplemental Figure 9). Taken together, our data indicate that IQGAP1 overexpression in tumors exacerbates Hippo/YAP signaling via enhanced NUAK2 expression, which may be a druggable mechanism for a specific subset of HCC patients.

## Discussion

Our results show that both loss and gain of IQGAP1, a scaffold protein, can promote HCC in the murine liver (Figure 7). Consistent with our finding, IQGAP1 is frequently induced and is associated with a worse prognosis in human HCC [11-14, 63]. Intriguingly, we found that IQGAP1-deletion also increased tumor burden. We reconcile these results and postulate that IQGAP1 has a bimodal effect on promoting hepatic tumorigenesis.

IQGAP1 loss does not predispose livers to develop differential subsets of liver cancer (e.g., HCA versus HCC) (Figure 1F) nor cause discrete molecular characteristics (Supplemental Figure 3). However, *Iqgap1-*deficient tumor lesions proliferate and rapidly develop liver cancer with increased severity (Figure 2D-E). We found that canonical Wnt/β-catenin signaling was unaltered in *Iqgap1*^*-/-*^ livers (Supplemental Figure 6), despite previous studies showing that IQGAP1 facilitates β-catenin’s nuclear translocation and activity [14, 43, 64]. Why β-catenin signaling does not change in the absence of IQGAP1 warrants future investigation. We found *in vitro* knock down of *IQGAP1* increased *CCND1* expression and MET expression in HepG2 cells (Figure 3B-D). MET expression was also induced in *Iqgap1*^*-/-*^ liver tumors suggesting that this increase may be compensatory to the loss of IQGAP1 expression, or IQGAP1 may regulate turnover of the MET protein.

On the other hand, we demonstrate that increased expression of IQGAP1 is sufficient to increase tumor burden and exacerbates HCC development in the B+M model (Figures 4C-E). This enhanced tumor burden is not driven by increased MET or Wnt signaling (Supplemental Figure 8 and Figure 5). Even though IQGAP1 overexpression can enhance β-catenin activity *in vitro* (Figures 5B-C) and significantly increase cytosolic β-catenin protein *in vivo* (Figure 5D), we find that β-catenin and IQGAP1 co-localize at the cell membrane in B+M+I livers. This implies that higher IQGAP1 prevents β-catenin nuclear translocation (Figure 5G) and is in line with a well-known mechanism regulated by IQGAP1 that enhances E-Cadherin-β-catenin complexes at the adherens junctions (Figures 5F-G) [44, 45, 65, 66].

HCC tumors are heterogenous and are not driven by a singular oncogenic pathway, and IQGAP1 is known to enhance multiple oncogenic pathways [28, 29, 67]. We found one such pathway in our model to be the Hippo/YAP signaling pathway. IQGAP1 is known to interact with and modulate YAP activity [48, 49]. We found significantly elevated expression of *Birc5*, a target strongly linked to YAP activity [47], when IQGAP1 is overexpressed in the B+M model (Figure 5E). In addition, we also found more YAP protein in the nucleus of B+M+I samples compared to B+M (Figure 6A) and its targets remained unchanged when comparing B+M to B+M+I samples (Figure 6B). This data matched well with recent findings that most YAP downstream proteins are unaltered in HCC except for NUAK2 (Figure 6C). To evaluate the translational relevance of this findings, we mined TCGA database. We found 7 patients with *IQGAP1*^*High*^/*NUAK2*^*High*^ expression (Figure 6D), who not only have increased proliferative signaling pathways but also exhibit poor survival (Supplemental Figure 8 and Figure 6E). Additionally, these patient samples have no activating *CTNNB1* mutations (data not shown), which indicates therapies targeting IQGAP1 and/or NUAK2 may be more beneficial. Globally, there were about 953,000 liver cancer cases in 2017 [68]. Considering that 1.9% of patients in the TCGA cohort have increased IQGAP1 and NUAK2 expression, this equates to roughly 18,000 patients per year worldwide. This might be an underestimate due to our use of TCGA mRNA expression data, since its known that 60-80% of HCCs display elevated IQGAP1 protein expression [11-14].

In summary, we show that overexpression and deletion of IQGAP1 promote hepatic tumorigenesis through two distinct mechanisms- by YAP signaling or by increased MET expression respectively. Such paradoxical findings have been reported in other key proteins involved in tumorigenesis, including NF-κB, JNK kinases, STAT3, MET, and β-catenin [69]. Our results indicate that IQGAP1 may function as a rheostat to balance pro-proliferative signals and that too little and too much may tip the balance towards uncontrolled proliferation. Therefore, targeting domain-specific interactions of IQGAP1 may be useful strategy to combat hepatic tumorigenesis.

## Methods

### Animals

The Institutional Care and Use Committee of approved all mouse experiments. *Iqgap1*^*+/+*^, *Iqgap1*^*+/-*^, and *Iqgap1*^*-/-*^ mice maintained on a 129/SVJ background (129-*Iqgap1*^*tm1aber*^*/*VsJ) were used for all diethylnitrosamine (DEN) experiments. These mice were generated in Dr. Andrew Bernards’s laboratory (Massachusetts General Hospital, Boston, Massachusetts, USA) and were obtained from Dr. Valentina Schmidt (Stony Brook University, New York, USA). FVB/NJ mice were obtained from Jackson Labs (Bar Harbor, ME). Animals were either housed at the University of Illinois at Urbana-Champaign on conventional racks or the University of Pittsburgh in Optimice cages (AnimalCare Systems, Centennial, CO) with Sani-Chip Coarse bedding (P. J. Murphy, Montville, NJ) at 24 °C on a 12/12 hour light/dark cycle with lights on starting at 5AM CST, corresponding to zeitgeber time (ZT) 0. Genotype was confirmed by PCR analysis of genomic DNA from tail clips. Mice were allowed *ad libitum* access to water and either Teklad F6 Rodent Diet (Envigo 8664) or standard mouse chow (LabDiet, St. Louis, MO, Purina ISO Pro Rodent 3000). Mice were provided huts and running wheels for enrichment. All animals were sacrificed between 9 am and noon daily.

### Mice experiments

Male (n = 77) and female (n =15) littermate *Iqgap1*^*+/+*^, *Iqgap1*^*+/-*^, and *Iqgap1*^*-/-*^ mice were injected with 5 mg/kg DEN (N0258, Sigma-Aldrich) in sterile 1x PBS at 12-15 days of age via intraperitoneal injection (10 µL/g body weight). Mice were sacrificed at both 20 weeks and 50 weeks after administration to assess tumor burden.

Six to eight-week-old male FVB/NJ mice were used for hydrodynamic tail vein injections. Mice were injected with 20 mg of pT3-EF5α-hMet-V5, pT3-EF5α -S45Y-β-catenin-Myc, or pT3-EF5α-IQGAP1-HA, or combination of EF5α-hMet-V5 and pT3-EF5α-S45Y-β-catenin-Myc, or combination of EF5α-hMet-V5, pT3-EF5α-S45Y-β-catenin-Myc and pT3-EF5α-IQGAP1-HA along with the sleeping beauty transposase (SB) (0.8 mg) in a ratio of 25:1. Injections were diluted into a total of 2 mL of normal saline (0.9% NaCl) and injected into the lateral tail vein in 5 to 7 seconds. See Supplemental Table 2 for construct information.

At the time of sacrifice, blood was collected by retro-orbital bleeding, and serum was separated by centrifugation and immediately stored at -80 °C in opaque tubes. Liver, gonadal white adipose tissue, spleen, and quadriceps tissues were collected, weighed, and flash frozen for analysis. A piece of each liver/tumor and the lungs were fixed in 10% formalin for histological analysis.

### Body weight and LW/BW

Livers from experimental animals were excised, washed in PBS and weighed. The percentage of the weight occupied by the liver was determined by dividing the liver weight by the total body weight of the mouse.

### Cell Lines

Human HepG2, Hep3B, Snu-449 and Huh7 hepatoma cell lines were obtained from American Type Cell Culture (ATCC). HepG2, Hep3B and Huh7 were maintained in 10% fetal bovine serum (FBS) (Atlanta Biologicals) in DMEM. Snu-449 cells were maintained in 10% FBS in RPMI-1640. Cells were incubated at 37°C in a humidified 5% carbon dioxide atmosphere.

### Constructs Used

pEGFP-IQGAP1 was a gift from David Sacks (Addgene plasmid # 30112). Using this construct, an HA tag was added to IQGAP1 and cloned via Gateway PCR (Invitrogen, Carlsbad, CA) into a pT3-EF5α vector. Additional constructs used can be found in Supplemental Table 2.

### Luciferase assay

Cell lines were transfected simultaneously with 400 ng TOPFlash firefly luciferase reporter and 100 ng Renilla luciferase constructs alongside either siRNA or expression constructs listed in Supplemental Table 2 using Lipofectamine 2000 (Life Technologies). Transfected cells were harvested after 72h and processed with Dual-Luciferase Reporter Assay kit (Promega). Luciferase activity was detected with an Infinite M200 PRO microplate reader (Tecan, Männedorf, Switzerland). Relative luciferase activities of transfected plasmids are represented as the activity of firefly luciferase activity normalized to Renilla activity.

### Lentivirus production and shRNA knock down

Lentiviruses packaged with shRNA were produced as previously described [70]. Constructs used are listed in Supplemental Table 2. HepG2 cells were infected with 25 µL of lentivirus-containing medium expressing either *shScrambled* or *shIQGAP1* constructs and infected cells were selected by culturing cells in 1µg/mL puromycin for 96 hours. Cell lysates were processed for RNA expression analysis or western blotting using TriZOL or RIPA buffer, respectively.

### Histology

Liver samples were fixed in formalin for > 24 hours. They were then processed and embedded in paraffin wax. Four or five µm sections were cut. For immunohistochemistry, sections were deparaffinized using xylene and graded ethanol (100-95%) washes and incubated in citric-acid based antigen retrieval (Vector, Burlingame, CA). Following antigen retrieval, liver sections were treated with 3% hydrogen peroxide to quench endogenous peroxidase and blocked with either 5% normal goat serum in 5% bovine serum albumin in TBST or Avidin/Biotin blocking solution (Vector SP-2001). Slides were incubated with primary antibody followed by biotinylated secondary antibody or HRP-conjugated secondary antibody (concentrations indicated in Supplemental Table 3). Either avidin-conjugated peroxidase (Vector ABC kit PK-6100) with ImmPACT DAB Peroxidase Substrate (Vector SK-4105) or DAB Peroxidase (HRP) Substrate Kit (SK-4100, Vector Labs) were used to visualize stained tissues. Sections were counterstained with Modified Harris Hematoxylin (Richard Allen 72711) dehydrated with ethanol and xylene washes and mounted with Permount (Fisher).

Briefly, H&E staining was performed after deparaffinization. Slides were first stained with hematoxylin and rinsed with water followed by dipping in 5% glacial acetic acid. Shandon’s Bluing Reagent (ThermoScientific, Kalamazoo, MI) was used to retain hematoxylin counterstain. Slides are then dipped in Eosin for 1 minute and dehydrated with ethanol and xylene washes followed by mounting with Permount.

### RNA isolation, quantitative reverse transcriptase polymerase chain reaction (qRT-PCR) and PCR

Total RNA from fresh liver and tumor samples collected at sacrifice was extracted using TRIzol solution (Invitrogen) and subjected to qRT-PCR to quantify the expression of protein-coding genes. A260/280 and bleach RNA gel were used to assess RNA quality. RNA with an A260/280 > 2.0 and a 28S/18S RNA ratio of approximately 2 was used for further analysis. Complementary DNA (cDNA) synthesis and qRT-PCR were performed either as previously described [24] or using M-MLV (Life Technologies) followed by q-PCR performed with SYBR Green PCR Master Mix (Life Technologies), Bullseye EvaGreen PCR Master Mix (Midwest Scientific), or Taqman probes (Life Technologies) with Taqman Universal Master Mix II (Life Technologies). Primer sequences and Taqman probe IDs are described in Supplemental Table 4. Reactions were performed using a StepOnePlus System (Life Technologies). Relative expression was calculated using the ΔΔCT method. *Gapdh* was used as housekeeping gene. Taqman probe IDs and primer sequences are listed below. Reactions were performed using a StepOnePlus System (Life Technologies).

### DNA mutation analysis

Genomic DNA was isolated from DEN-induced liver tumors at 50 weeks post-treatment using QIAamp Fast DNA tissue kit (Qiagen). Fludigm technology was used to sequence the DNA using four primer sets (Supplemental table 4). The read files were demultiplexed by primer and then by sample. Fastp version 0.19.5 was used to perform quality filtering and trimming of the raw reads. The files were qc-trimmed and aligned to the mouse reference genome version GRCm38.p6 with NCBI WebBlast in order to obtain the absolute coordinates in the forward strand version of the genome with bwa version 0.7.17 using default parameters. Bam-read count version 0.8 was run to generate the read counts at each of the locations identified. Then a customized R script was run to generate a summary of nucleotide frequencies at the codons of interest. The base pair with the lowest percent of reads matching the WT sequence was used to determine the prevalence of mutation at that codon in each sample. The raw sequence data have been uploaded to NCBI Bioproject (http://www.ncbi.nlm.nih.gov/bioproject/629000) and will be made public after publication.

### Western blotting

Whole tissue protein lysates were prepared from approximately 50 mg frozen tissue using sodium dodecyl sulfate (SDS)-based lysis buffer (50 mM Tris-HCl, pH 8.0, 10 mM EDTA, 1% SDS) containing protease/phosphatase inhibitors (RIPA buffer). Lysates were removed to a fresh 1.5 ml tube and centrifuged at 18.4g for 10 min. at 4°C in order to remove clear supernatant to a new 1.5 ml tube while disposing of the pellet. Samples were stored at - 80°C until utilization or determination of protein concentration via BCA protein assay (Fisher) to ensure equal protein concentrations for subsequent assays. For Western blot, 50-200 µg total protein was loaded onto 8%-12% SDS-PAGE gels. Protein were transferred to Immobilon-P PVDF membrane (IPVH00010, Millipore) either for 1 hour at 100V at 4°C or overnight at 35V and 4°C. After transfer, the membranes were blocked in either 5% non-fat dry milk or 5% BSA dissolved in Blotto (0.15M NaCl, 0.02M Tris pH 7.5, 0.1% Tween in dH2O) followed by incubation with antibodies described in Supplemental Table 3. Membranes were exposed to SuperSignal West Pico Chemiluminescent Substrate (Thermo Scientific Pierce, Pittsburgh, PA) for 1-2 min. at room temperature and bands reflective of target proteins were viewed by ChemiDoc imaging system (Bio-Rad). Bands were quantified with ImageJ (National Institutes of Health, Bethesda, MD).

### Nuclear/Cytoplasmic Fractioning

Whole liver samples were processed either by using the NE-PER Nuclear and Cytoplasmic Extraction kit (Life Technologies) following manufacturers recommendations or lysed using SDS-free subcellular fractionation buffer (20 mM HEPES, 10 mM KCl, 2 mM MgCl_2_, 1 mM EDTA, 1 mM EGTA) containing protease/phosphatase inhibitors. Samples were manually agitated and incubated on ice for 30 minutes prior to centrifugation at 3000 RPM for 5 minutes at 4°C. The supernatant containing the cytosolic fraction was moved to a fresh, 1.5 ml tube while the pellet was washed 5x with subcellular fractionation buffer. After final wash, nuclear pellet was lysed using TBS with 0.1% SDS. Samples were stored at -80°C until use in protein quantification and western blotting as previously described.

### Immunoprecipitation

Whole liver samples were lysed using immunoprecipitation lysis buffer (20 mM Tris-base, 150 mM NaCl, 1 mM EDTA, 1 mM EGTA, 1% Triton-X pH 7.5) containing protease/phosphatase inhibitors. Samples were manually agitated and incubated on ice for 30 minutes prior to centrifugation at 15,000 RPM for 15 minutes at 4°C. Supernatants were removed to a fresh 1.5 ml tube while the pellet was discarded. Samples were stored at -80°C until utilization or determination of protein concentration via BCA protein assay (Fisher) to ensure equal protein concentrations. Equal amounts of protein from respective groups were pooled together and precleared on ice using normal control mouse IgG (Life Technologies) for 30 minutes followed by incubating with A/G PLUS-Agarose beads (Santa Cruz) overnight at 4°C with gentle agitation. Samples were centrifuged for 5 minutes at 3000 RPM and supernatants were removed to a fresh 1.5 ml tube together with 5 µg of β-catenin monoclonal antibody (Supplemental Table 3) overnight at 4°C with gentle agitation. A/G PLUS-Agarose beads were applied to each sample overnight at 4°C with gentle agitation. The supernatant was then removed from beads via centrifugation to a fresh 1.5 ml tube and the pellet was washed four times with PBS containing protease/phosphatase inhibitor to prevent disruption of delicate protein-protein interactions. Samples were then processed for western blotting as previously described.

### Data Analysis

RNA-Sequencing gene expression data from the Hepatocellular Carcinoma TCGA Firehose Legacy dataset was downloaded from the cbioportal (cbioportal.org) [71, 72]. Patients were separated into two cohorts, those that have amplified *IQGAP1* and *NUAK2* mRNA expression fitting a cutoff z-score threshold ± 2, and those without. Gene expression data were analyzed through the use of IPA (QIAGEN Inc., https://www.qiagenbioinformatics.com/products/ingenuitypathway-analysis) using an experimental p-value > 0.02 and a false discovery rate q-value > 0.04 [73]. Canonical pathway amplification/downregulation was determined using a -log(p-value) < 2.3 and a z-score < 2.5 while thresholding at 0.05. Fisher’s exact test was used to determine significance of pathway alterations. Finally, pathways were filtered for relevance to liver biology and disease pathogenesis. For survival analysis, clinical data from the TCGA were analyzed to determine overall survival calculated from diagnosis date to the death date or date of last contact taking censoring into consideration. Overall survival was then calculated for the patients using Kaplan-Meier methods.

### Statistical Analysis

Data are expressed as mean ± SD or SEM. All statistical analyses were performed using GraphPad Prism software version 7. For contingency data, χ^2^ test was used to compare 3 groups and Fisher’s exact test was performed to assess differences between 2 groups. One-way ANOVA with Bonferroni post-test was performed to compare 3 groups while two-way ANOVA with Tukey’s post-test was used to assess differences between two paired tissues (liver and tumor) in three groups. Asterisks indicate a statistically significant difference between groups. Significance is defined as P < 0.05. Outliers were determined by Grubbs’ test and were removed from analysis along with any paired data.

### Study Approval

Animal studies were approved by the Institutional Animal Care and Use Committees at the University of Illinois at Urbana-Champaign and University of Pittsburgh. All animal studies were carried out as outlined in the *Guide for the Care and Use of Laboratory Animals* [74].

## Conflict of interest disclosure statement

There are no conflicts of interest.

## Acknowledgements

IPA licensed through the Molecular Biology Information Service of the Health Sciences Library System, University of Pittsburgh was used for data analysis.

## Authors’ Contributions

Writing, review, and/or revision of the manuscript: H.L.E., E.R.D., A.W.D., S.A.

Technical, or material support (experimental design, execution, data and statistical analysis): .L.E., E.R.D., J.T, S.P.M., A.W.D, S.A. Study supervision: A.W.D., S.A.

## Financial support

S.A.: R01 DK113080, ACS132640-RSG

A.W.D.: R01 DK103645

H.L.E.: F30 CA206495

## Notes

### Competing Interest Statement

The authors have declared no competing interest.

