## Supplemental data for "Deletion and overexpression of the scaffolding protein IQGAP1 promotes HCC"

**Supplemental Table 1 Molecular Subtype Breakdown of HCC Tumors**

| G1/2 | G3 | G4 | G5/6 |
|---|---|---|---|
| | *Rrm2* | *Rrm2* | *Rrm2* |
| | *Angpt2* | *Angpt2* | |
| *Glul* | | *Glul* | *Glul* |
| | *Tgfbr1* | *Tgfbr1* | |
| | | *Crp* | *Crp* |
| | *Pepck* | *Pepck* | *Pepck* |
| *Fasn* | *Fasn* | *Fasn* | |

Gene expression patterns correlating with molecular subtypes of HCC adapted from Boyault et al. 2007. Red represents upregulated genes. Blue represents downregulated genes.

**Supplemental Table 2 Constructs and siRNA used.**

| *Constructs/siRNA* | *Species* | *Use* | *Source* |
|---|---|---|---|
| pT3-EF5α-hMet-V5 | Human | *In vivo* | Monga Lab |
| pT3-EF5α -S45Y-β-catenin-Myc | Human | *In vivo* | Monga Lab |
| pT3-EF5α-IQGAP1-HA | Human | *In vivo* | Custom |
| pCMV/Sleeping Beauty Transposase | N/A | *In vivo* | Monga Lab |
| pIRES2-EGFP | N/A | *In vitro* | Clonetech |
| pEGFP-IQGAP1 | Human | *In vitro* | Courtesy of Dr. David Sacks (Addgene) |
| S45Y-β-catenin | Human | *In vitro* | Monga Lab |
| TOPFlash Firefly Luciferase Reporter | N/A | *In vitro* | Monga Lab |
| Renilla Luciferase | N/A | *In vitro* | Monga Lab |
| *si-Scrambled Control* | N/A | *In vitro* | Life Technologies Cat# 12935-300 |
| *si-IQGAP1* | Human | *In vitro* | Life Technologies Cat# s16837 |
| *si-CTNNB1* | Human | *In vitro* | Life Technologies Cat# 42816 |
| *sh-Scrambled* | N/A | *In vitro* | Courtesy of Dr. Jie Chen Cat# 1864 (Addgene) |
| *sh-IQGAP1* | human | *In vitro* | Sigma Aldrich Cat# TRCN0000047485 |

**Supplemental Table 3 Antibodies.**

| *Antibodies and associated reagents* | *Vendor* | *Catalog #* | *Dilution* | *Use* |
|---|---|---|---|---|
| 1234/1235-MET | Cell Signaling | 3077 | 1:1000 | Western Blot |
| AKT-1 | Santa Cruz | sc-5298 | 1:1000 | Western Blot |
| β-catenin | Santa Cruz | sc-7963 | 1:1000 | Western Blot |
| E-Cadherin | BD Biosciences | 610181 | 1:1000 | Western Blot |
| GAPDH | Santa Cruz | sc-25778 | 1:1000 | Western Blot |
| HA | Santa Cruz | sc-7392 | 1:200 | Western Blot |
| IQGAP1 | Santa Cruz | sc-10792 | 1:1000 | Western Blot |
| LaminB1 | Santa Cruz | sc-374015 | 1:100 | Western Blot |
| MET | Cell Signaling | 3127 | 1:500 | Western Blot |
| mTOR | Santa Cruz | sc-517464 | 1:200 | Western Blot |
| MYC | Santa Cruz | sc-40 | 1:200 | Western Blot |
| S2448-mTOR | Santa Cruz | sc-293133 | 1:1000 | Western Blot |
| S473-AKT-1/2/3 | Santa Cruz | sc-514032 | 1:350 | Western Blot |
| STAT3 | Santa Cruz | sc-8019 | 1:500 | Western Blot |
| V5 | eBioscience | 14-6796-82 | 1:200 | Western Blot |
| Y705-STAT3 | Santa Cruz | sc-8059 | 1:1000 | Western Blot |
| YAP1 | Santa Cruz | sc-101199 | 1:200 | Western Blot |
| Donkey anti-Rabbit/Horseradish peroxidase | GE Healthcare Life Sciences | NA934V | 1:5,000 | Western Blot |
| Goat anti-Mouse/Hoseradish peroxidase | Invitrogen | PI31444 | 1:25,000 | Western Blot |
| Glutamine Synthetase | Abcam | ab49873 | 1:300 | Immunohistochemistry |
| P-ERK1 | Cell Signaling | 4370 | 1:400 | Immunohistochemistry |
| Ki67 | BD Biosciences | 550609 | 1:250 | Immunohistochemistry |
| Biotinylated goat anti-rabbit antibody | Vector Laboratories | BA-1000 | 1:1000 | Immunohistochemistry |
| Goat anti-rabbit HRP | GE Healthcare | NA9340 | 1:250 | Immunohistochemistry |
| Goat anti-mouse HRP | Bio-Rad | 170-6516 | 1:250 | Immunohistochemistry |

**Supplemental Table 4 Primers and Taqman Probes**

| Gene | Species | Forward Primer (5' to 3') | Reverse Primer(5' to 3') |
|---|---|---|---|
| *AXIN2* | *Human* | GCTCCAGAAGATCACAAAGAGC | AGCTTTGAGCCTTCAGCATC |
| *B2M* | *Human* | TGAGTATGCCTGCCGTGTG | ATTCATCCAATCCAAATGCGG |
| *CCND1* | *Human* | TCAAGACGGAGGAGACCTGT | GGAAGCGGTCCAGGTAGTTC |
| *CTNNB1* | *Human* | GAAACGGCTTTCAGTTGAGC | CTGGCCATATCCACCAGAGT |
| *GAPDH* | *Human* | ATGACATCAAGAAGGTGGTGAA | GCTGTTGAAGTCAGAGGAGAC |
| *IQGAP1* | *Human* | AGGCAAGCAAACTGCCCTAT | CTGATGGAGCTGTCTAGCCG |
| *MET* | *Human* | TTACGGACCCAATCATGAGC | GCTGCAAAGCTGTGGTAAAC |
| *Afp* | *Mouse* | CCGGAAGCCACCGAGGAGGA | TGGGACAGAGGCCGGAGCAG |
| *Angpt2* | *Mouse* | ACAGCTGTGATGATAGAGATTGG | CGAGTCTTGTCGTCTGGTTTAG |
| *Axin2* | *Mouse* | GCTCCAGAAGATCACAAAGAGC | AGCTTTGAGCCTTCAGCATC |
| *Birc5* | *Mouse* | TTTCCAAATACCACTGTCTCCTTCTC | GCCACGCATCCCAGCTT |
| *Crp* | *Mouse* | CGTATGGCGGTGACTTTGA | GTGCTGATCTGTTCTGGAGATAG |
| *Ctnnb1* | *Mouse* | ACTTGCCACACGTGCAATTC | AAGGTTGTGCAGAGTCCCAG |
| *Ccnd1* | *Mouse* | TCAAGACGGAGGAGACCTGT | GGAAGCGGTCCAGGTAGTTC |
| *Fasn* | *Mouse* | GCTGCGGAAACTTCAGGAAAT | AGAGACGTGTCACTCCTGGACTT |
| *Gapdh* | *Mouse* | TCCTGCACCACCAACTGCTTAG | TGCTTCACCACCTTCTTGATGTC |
| *Glul* | *Mouse* | CAGGCTGCCATACCAACTTCA | TCCTCAATGCACTTCAGACCAT |
| *Iqgap1* | *Mouse* | GATTCCCTGCACGAGAAGTT | CGATGGCTGGGTTCATGTAT |
| *Iqgap2* | *Mouse* | TCAAGATTGGACTGCTGGTG | AGGTTTGGTCTGGAGGAGGT |
| *Iqgap3* | *Mouse* | CTCTGGTCACCTTGCAGAAT | CAGCAGCTCTTGGTAGACAG |
| *Nuak2* | *Mouse* | CAGGCATTTCTTCCGACAGATC | GAGAGGCCAAAGTCAGCAAT |
| *Lect2* | *Mouse* | CCCACAACAATCCTCATTTCAGC | ACACCTGGGTGATGCCTTTG |
| *Pepck* | *Mouse* | GTTCCCAGGGTGCATGAAAG | AGGGCGAGTCTGTCAGTTCAA |
| *Rrm2* | *Mouse* | GTTGTCTTTCCCATCGAGTACC | GAGCTTCCCAGTGCTGAATATC |
| *Tgfbr1* | *Mouse* | GGGCTTAGTGTTCTGGGAAA | CCGATGGATCAGAAGGTACAAG |
| *Egfr F254* | *Mouse* | CATGTTGTCCCTCCTATGACAG | GGGCACTTCTTCACACAGGT |
| *Braf V537* | *Mouse* | TTCCTTTACTTACTGCACCTCAGA | CCTCCTCAATTCTTACCATCCA |
| *Braf K584* | *Mouse* | CTTTTTGGCAGGAAAACTCG | ACTTACTCCATGCCCTGTGC |
| *Hras Q61* | *Mouse* | CCCACTAAGCCGTGTTGTTTT | TCCTCGAAGGACTTGGTGTT |
| *Amotl2* | *Mouse* | *Taqman Probe ID#* | Mm00502287_m1 |
| *Ccn1* | *Mouse* | *Taqman Probe ID#* | Mm01323719_g1 |
| *Ccn2* | *Mouse* | *Taqman Probe ID#* | Mm01192933_g1 |
| *Gapdh* | *Mouse* | *Taqman Probe ID#* | Mm99999915_g1 |
| *Jag1* | *Mouse* | *Taqman Probe ID#* | Mm00496902_m1 |
| *Yap1* | *Mouse* | *Taqman Probe ID#* | Mm01143263_m1 |

**Supplemental Figure 1**

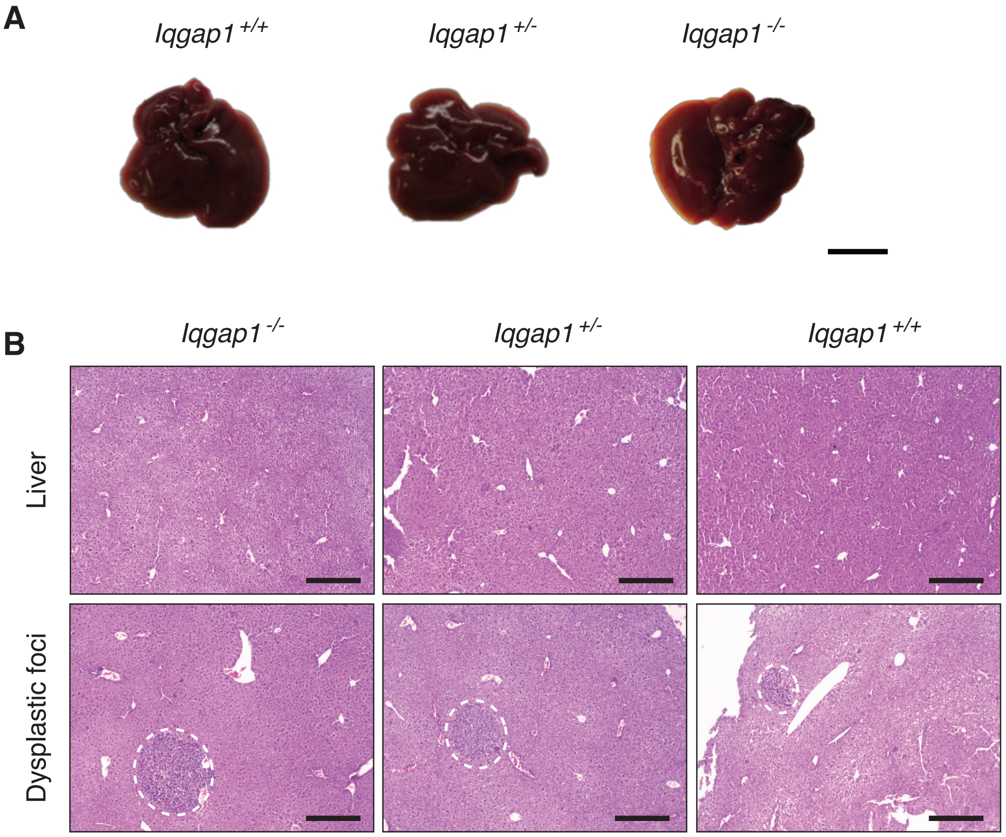


**Supplemental Figure 1. Dysplastic foci are observed 20 weeks post-DEN treatment.** (A) Representative liver photos from mice 20 weeks after DEN treatment (n = 9, 10, 5 *Iqgap1*$^{+/+}$, *Iqgap1*$^{+/-}$, and *Iqgap1*$^{-/-}$ mice per group, respectively). Scale bar is 1 cm. (B) Representative H&E images of livers from *Iqgap1*$^{+/+}$, *Iqgap1*$^{+/-}$, and *Iqgap1*$^{-/-}$ mice 20 weeks post-DEN treatment. Dashed lines indicate dysplastic regions. Scale bar is 100 $\mu$m.

### Supplemental Figure 2

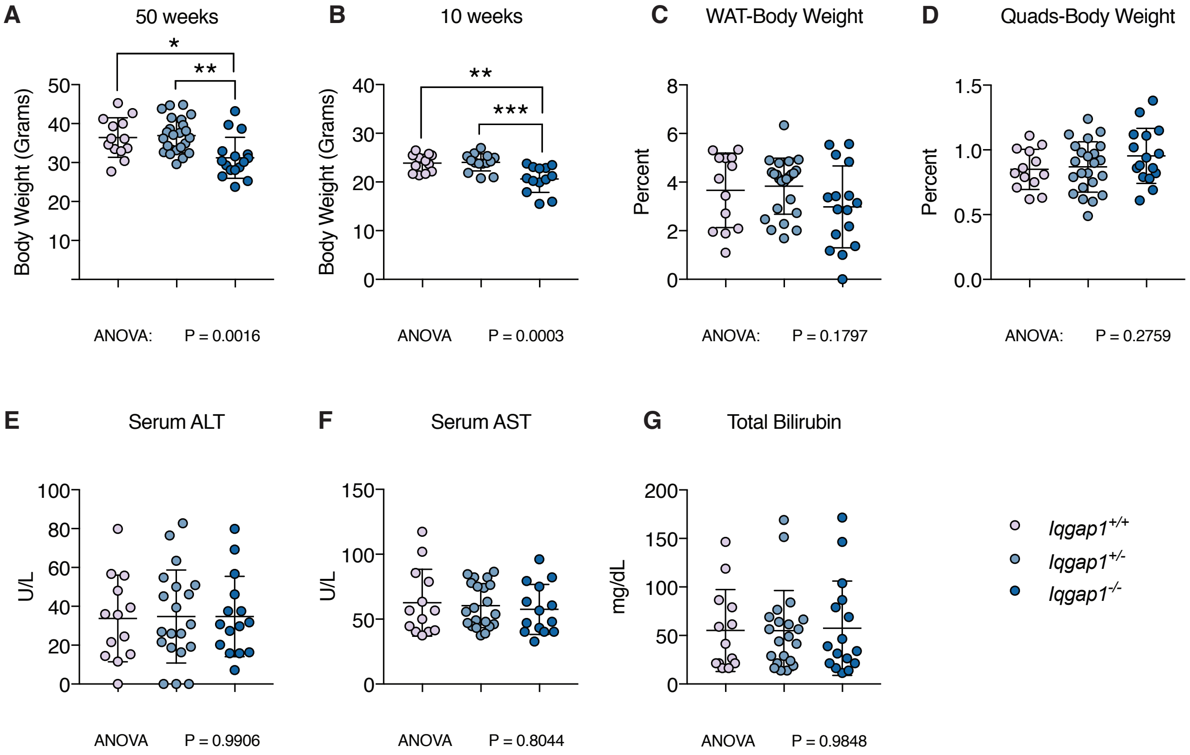


**Supplemental Figure 2. Characterization of DEN-treated mice at 50 weeks.** Male *Iqgap1*$^{+/+}$ (n = 13), *Iqgap1*$^{+/-}$ (n = 24), and *Iqgap1*$^{-/-}$ (n = 16) mice were treated with 5 mg/kg DEN at 12-15 days of age. After 50 weeks, tissues were harvested for analysis. (A) Body weight at time of sacrifice. (B) Body weight of DEN-treated *Iqgap1*$^{+/+}$, *Iqgap1*$^{+/-}$, and *Iqgap1*$^{-/-}$ mice 10 weeks after DEN treatment. (C) Gonadal white adipose tissue (WAT) weight normalized to total body weight. (D) Quadriceps muscle (Quads) weight normalized to body weight. (E-G) Serum levels of ALT (E), AST (F), and total bilirubin (G) were measured. Hemolyzed samples were excluded from analysis. Values are displayed as mean ± SD. One-way ANOVA with Bonferroni's multiple comparisons test was used to determine significance between groups. Significance is indicated by * $P < 0.05$, ** $P < 0.01$, *** $P < 0.001$.

### Supplemental Figure 3

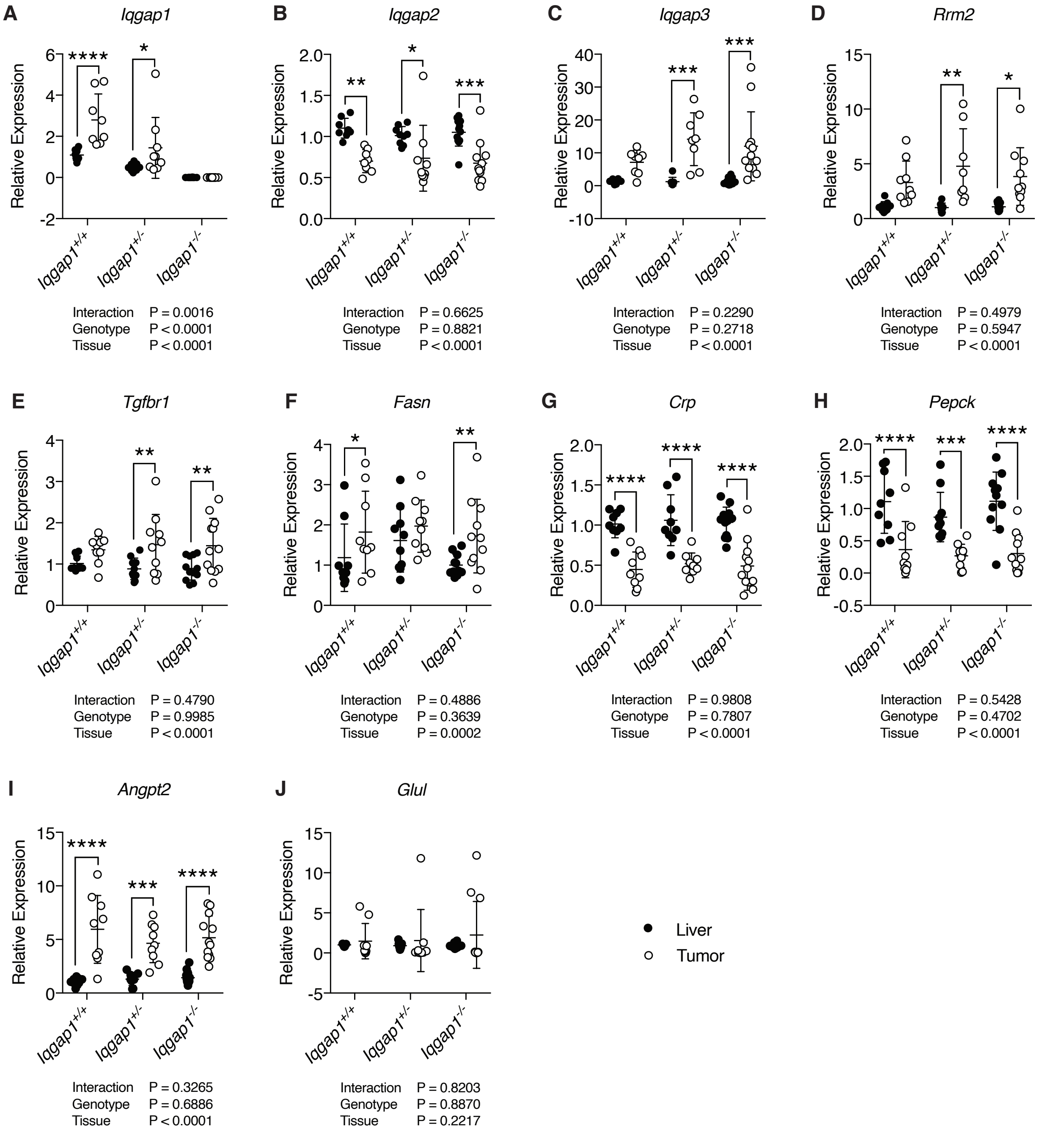


**Supplemental Figure 3. Hepatic gene expression in DEN tumors.** Gene expression of (A) *Iqgap1,* (B) *Iqgap2,* (C) *Iqgap3,* (D) *Rrm2*, (E) *Tgfbr1,* (F) *Fasn,* (G) *Crp,* (H) *Pepck,* (I) *Angpt2,* and (J) *Glul* in tumor-adjacent liver tissue and tumor tissue normalized to *Gapdh* expression. Values are displayed as mean ± SD. Two-way ANOVA with Tukey's multiple comparisons test was used to determine significance. Significance is indicated with * P < 0.05, ** P < 0.01, *** P < 0.001, **** P < 0.0001.

### Supplemental Figure 4

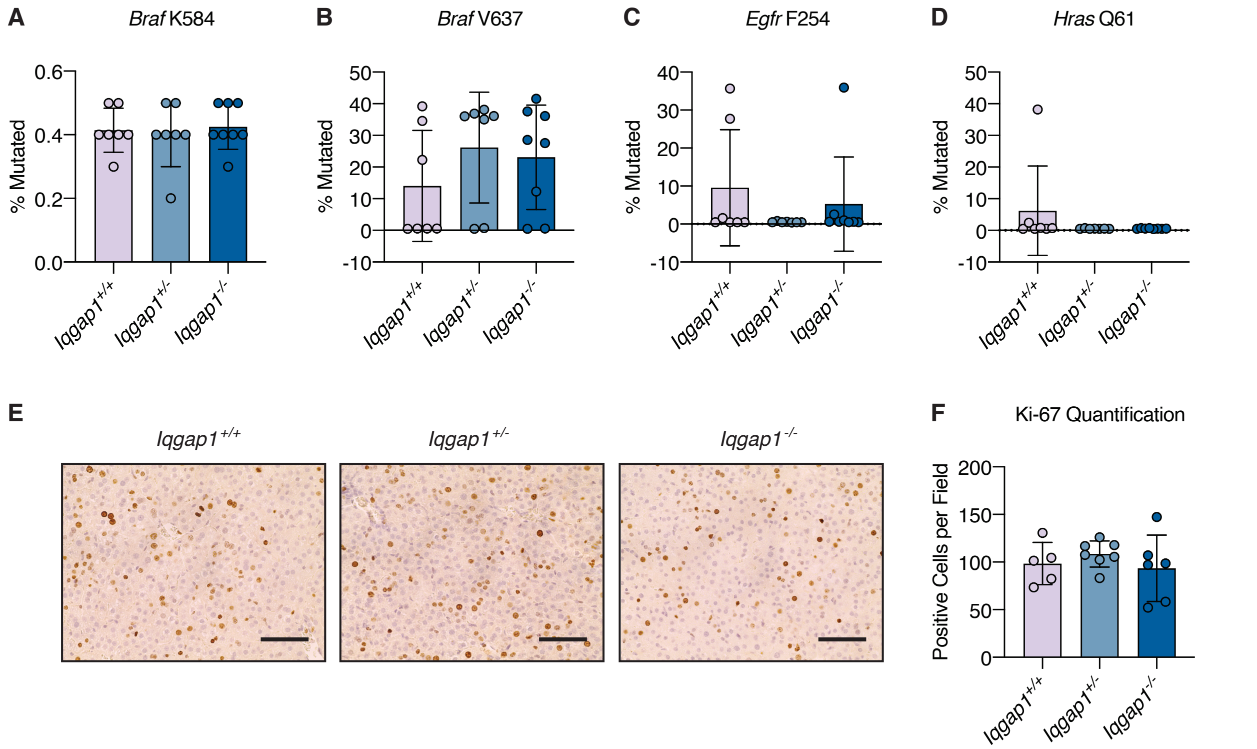


**Supplemental figure 4. IQGAP1 deletion does not affect DEN response.** (A-D) DNA mutation frequency at select codons in tumors of mice 50 weeks-post DEN injection (n = 7 *Iqgap1*$^{+/+}$, 7 *Iqgap1*$^{+/-}$, and 8 *Iqgap1*$^{-/-}$ mice). (E) Representative Ki-67 immunohistochemistry images of livers of P15 mice 24 hours after DEN injection. Scale bar is 50 $\mu$m. (F) Quantification of Ki-67-positive cells per field (n = 5 *Iqgap1*$^{+/+}$, 7 *Iqgap1*$^{+/-}$, and 6 *Iqgap1*$^{-/-}$ mice). One-way ANOVA with Bonferroni multiple comparisons test was used to compare groups.

**Supplemental Figure 5**

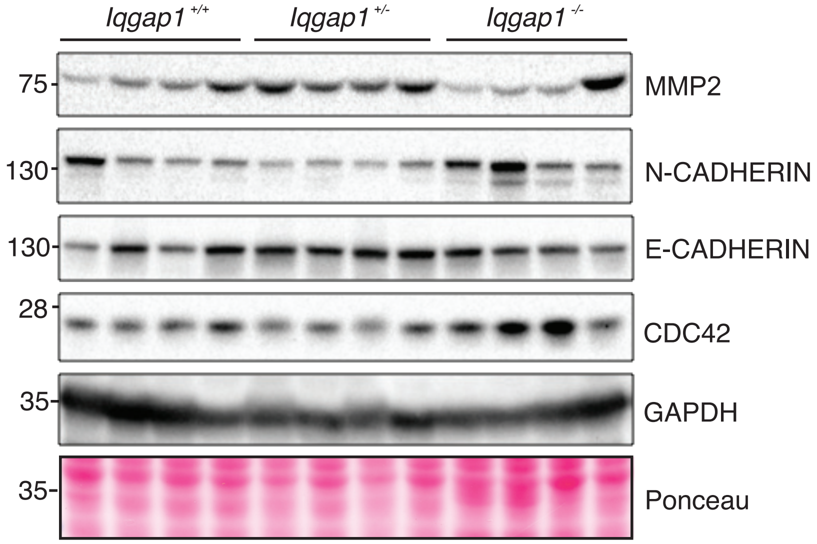


**Supplemental Figure 5. EMT is not affected by IQGAP1 deletion.** Immunoblot of EMT markers MMP2, N-Cadherin, E-Cadherin, and Cdc42 in tumors of *Iqgap1*$^{+/+}$*, Iqgap1*$^{+/-}$, and *Iqgap1*$^{-/-}$ mice 50 weeks after DEN-treatment (n = 4 mice per group).

**Supplemental Figure 6**

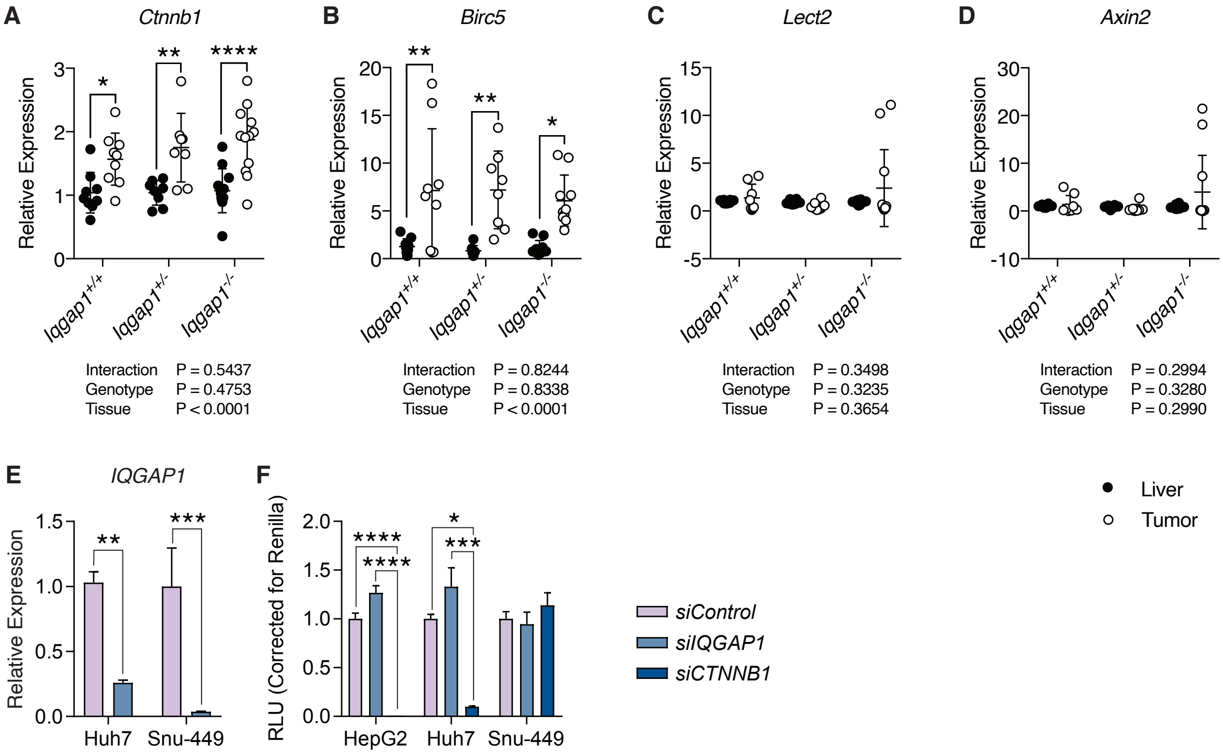


**Supplemental Figure 6. IQGAP1-deletion does not affect β-catenin activity.** Gene expression of (A) *Ctnnb1,* (B) *Birc5,* (C) *Lect2,* and (D) *Axin2* in tumor-adjacent liver tissue and tumor tissue normalized to *Gapdh* expression. (E) Confirmation of IQGAP1 knock down in Huh7 and Snu-449 cells. (F) Relative luciferase unit (RLU) measured from TOPFlash luciferase reporter assay in HepG2, Huh7, and Snu-449 cells transfected with *siControl*, *siIQGAP1*, and *siCTNNB1*. Values are displayed as mean ± SD. Two-way ANOVA with Tukey's multiple comparisons test was used to determine significance. Significance is indicated with * P < 0.05, ** P < 0.01, *** P < 0.001, **** P < 0.0001.

**Supplemental Figure 7**

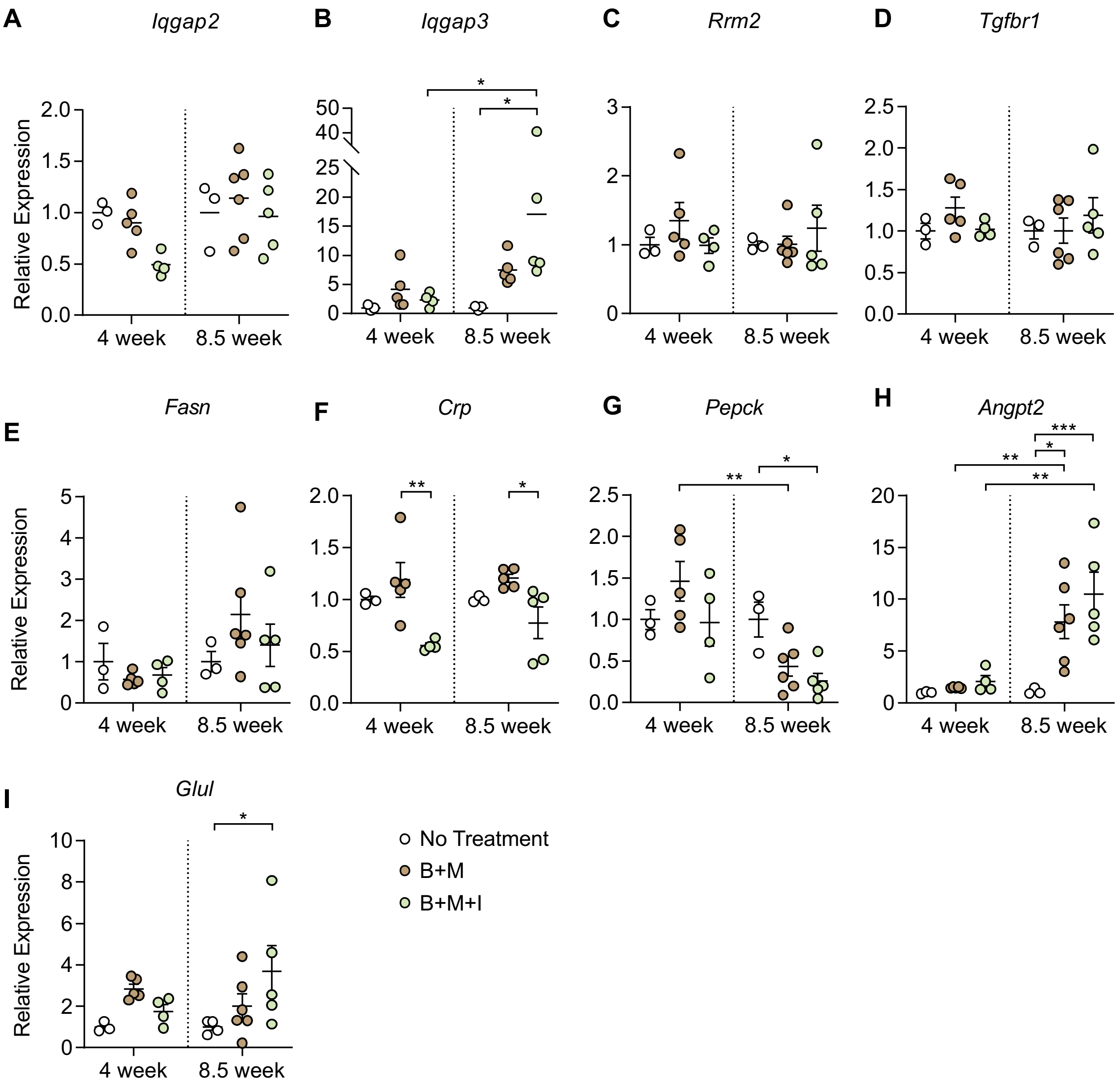


**Supplemental Figure 7. Hepatic gene expression in tumors induced via transposon system**. Gene expression of (A) *Iqgap2,* (B) *Iqgap3,* (C) *Rrm2,* (D) *Tgfbr1,* (E) *Fasn,* (F) *Crp,* (G) *Pepck,* (H) *Angpt2*, and (I) *Glul* in whole livers from 4- and 8.5-week samples from the transposon model. Gene expression was normalized to *Gapdh*. Values are displayed as mean ± SEM and dots represent individual mice. Two-way paired ANOVA with Tukey's multiple comparisons test was used to determine significance. Significance is indicated with * P < 0.05, ** P < 0.01, *** P < 0.001.

**Supplemental Figure 8**

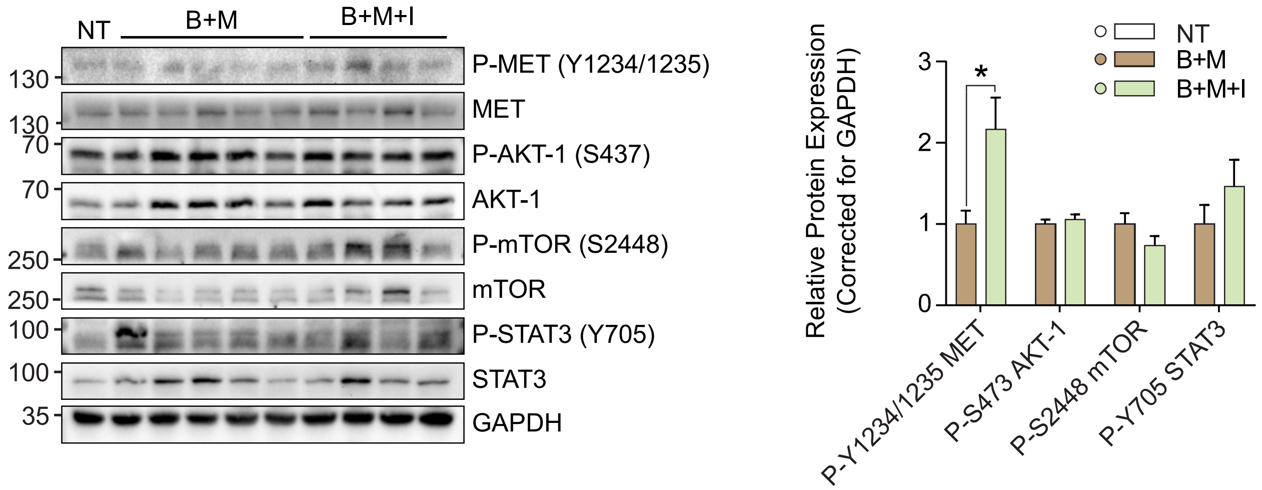


**Supplemental Figure 8. Increased IQGAP1 expression does not promote MET signaling *in vivo*.** Whole liver protein lysates analyzed by immunoblot for total and phosphorylated tyrosine residues of MET (Y1234/1235), AKT-1 (S437), mTOR (S2448), and STAT3 (Y705). Phosphorylated protein is normalized to GAPDH and corrected for total respective protein; values are expressed relative to NT control (set to 1). Graph show mean ± SEM. Significance is indicated with * $P < 0.05$

**Supplemental Figure 9**

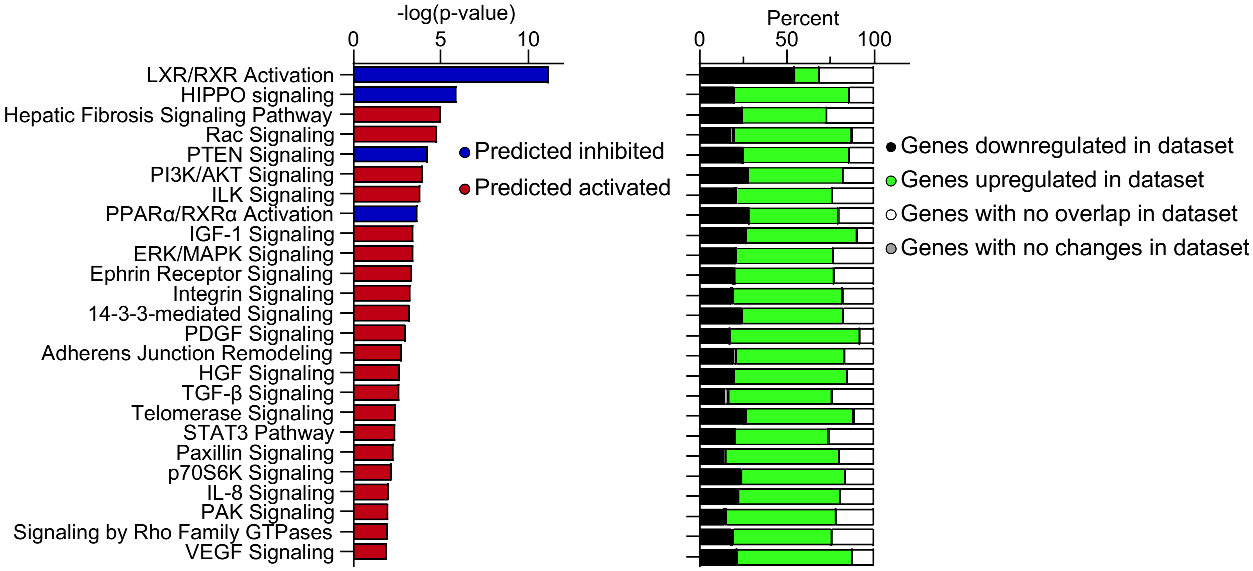


**Supplemental Figure 9. Molecular profiles of HCCs with *IQGAP1*$^{High}$/*NUAK2*$^{High}$ expression.** RNA-Sequencing data from the TCGA samples with elevated *IQGAP1*/*NUAK2* expression compared to those without was used for IPA analysis to determine molecular pathways that are activated/inhibited (left) with corresponding changes to the genes that regulate respective molecular pathways (right).
